# Measuring morphogen transport over multiple spatial scales in live zebrafish embryos

**DOI:** 10.1101/2025.02.06.636788

**Authors:** Ashwin V. S. Nelanuthala, Bitan Saha, Jagadish Sankaran, Tom J. Carney, Karuna Sampath, Thorsten Wohland

## Abstract

Morphogenesis is controlled by signalling morphogens that form gradients across the embryo. The gradient formation requires that the low molecular weight morphogens slow down their diffusion by at least one order of magnitude. However, the precise slow-down of diffusion across the relevant micrometre scales has not been directly observed. Here, we develop and employ Single-Plane Illumination Microscope based spatial Fluorescence Cross-Correlation Spectroscopy (SPIM-sFCCS) to directly measure the diffusion coefficient of the morphogen Squint in early zebrafish embryos as a function of topography and length scale. We show that Squint’s diffusion coefficient changes on length scales that are commensurate with the diameter of the intercellular spaces in the embryo and that the slow-down is regulated by receptor binding. The slowdown can be reduced by either the knockdown of Activin receptor 2b, a receptor for Squint, or the overexpression of Lefty2, an inhibitor of Squint. Based on our results and supporting simulations, we propose an interstitial space-dependent transient receptor binding and diffusing mechanism to explain this slowdown, which is crucial for gradient formation and embryonic patterning.

## 1 Introduction

At the heart of morphogenesis are concentration gradients of morphogens that control cell fates and patterns [1], [2]. The movement of morphogens is regulated by their interactions with cell surface receptors [3] and the extracellular matrix [4]. These interactions trigger the expression of key downstream genes, activators, inhibitors, and degradation pathways, all of which shape the morphogen gradient over space. The mobility of these signalling molecules is thus a limiting factor that dictates the rate at which signals propagate over space in an embryo and when and where biochemical signalling pathways and developmental processes are carried out. Therefore, maintaining robust transport dynamics of morphogens across space is vital for normal developmental events to take place at the right place and time.

To convey specific positional information to different cellular locations, morphogens are transported across space either through passive diffusion [5], active transport (transcytosis [6], [7], [8] or cytonemes [9]. In the case of diffusion, concentration gradients are formed by the interplay between diffusion and the degradation rate of the ligand. The gradient length is 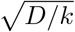, where D is the diffusion coefficient, and k is the degradation constant of the ligand. However, measured degradation rates, as far as available, are low (β 10^†4^/s) [10], [11], [12] and morphogens are signalling ligands of low molecular weight and, accordingly, have high D values (10-100 *μm*^2^*/s*). These parameters would result in an ineffective gradient length much larger than the embryo. Accordingly, global morphogen diffusion coefficients (*D*_*glob*_) measured over a large region (*>* 100 *μm*) are lower by one to two orders of magnitude compared to their local diffusion coefficient (*D*_*loc*_). This slowing down of morphogens from the local to global scale is seen in several morphogens, including Nodals, Wnts, BMP, and FGF [13] and references therein) and is crucial for the establishment of robust concentration gradients over space [14].

Several factors that cause ligand mobility to slow down have been demonstrated *in vivo*. Tortuosity, i.e. the hindrance of movement of particles due to the presence of cells, has been shown to slow down molecules in tissues [15], [16] and references therein. Reduced mobility by either binding to the extracellular matrix elements or being physically hindered by their presence has also been shown to contribute to the slowdown of morphogens [17], [18]. Moreover, transient receptor binding has been demonstrated to hinder morphogen movement by slowing down its effective mobility across space [12], [18]. In fact, it was demonstrated that tuning receptor concentrations in synthetic ligand-receptor pairs can influence the global diffusivity of ligands [19] and that receptor binding-based hindrance was sufficient to form synthetic concentration gradients of ligands *in vivo* [20]. Additionally, the geometry of extracellular space regulating the binding propensity of ligands to receptors was also demonstrated using single particle tracking *in vivo* [21].

Although the reasons for the slowdown are well established, the extent to which each factor contributes to the slowdown and the exact mechanism of how these factors combine to hinder diffusion need to be adequately addressed. Moreover, the length scales at which each of these factors operate to produce the transition from a fast to a slow diffusion coefficient need to be established. Probing the rate and scale of the transition from molecular size-dependent mobility to spatial interaction-hindered mobility allows a greater understanding of the rate of signalling processes that occur at characteristic length scales and helps model developmental processes.

To understand the extent and the length scale at which the slowdown occurs, we need to map the transition of morphogen diffusion from the local to the global scale. However, the commonly used *in vivo* morphogen diffusion measurement techniques do not cover all spatial scales. The *D*_*glob*_ measured using Fluorescence Recovery After Photobleaching (FRAP) [10], [22], [23] quantifies the recovery rate of fluorescence caused by diffusing fluorescent molecules into a large photobleached region of interest (10-100 *μm*). It thus is affected by the tissue morphology or tortuosity, and also by interactions of the morphogens with components of the extracellular space and cells (Fig. 1A, B). *D*_*loc*_, on the other hand, is measured by single molecule sensitive techniques. For example, Fluorescence Correlation Spectroscopy (FCS) *in vivo* [17], [24], [25], quantifies the time-span of the fluctuation of fluorescence signals arising from molecules diffusing in and out of a confocal volume, which is limited to *<* 1 *μm*. It thus recovers the unencumbered free diffusion coefficient determined only by particle size, viscosity, temperature, and possibly interactions with the extracellular matrix (Fig. 1C). Single Particle Tracking (SPT) follows single molecules in their trajectory in time with excellent spatial resolution. It thus could, in principle follow single molecules on their trajectory. However, due to the limited photostability of fluorophores, the single molecule trajectories rarely exceed several microns. Neither of these techniques can fully bridge the gap between the local and global scale and thus the intermediate length scales (1-10 *μm*) are hard to probe. Therefore, to determine the scale of influence of these diffusion regulators and probe the scale-dependent transition, there is a need to bridge the spatial information gap at the mesoscale (1-10 *μm*) between the local FCS and the global FRAP measurements (Fig. 1B, D).

**Fig. 1.**
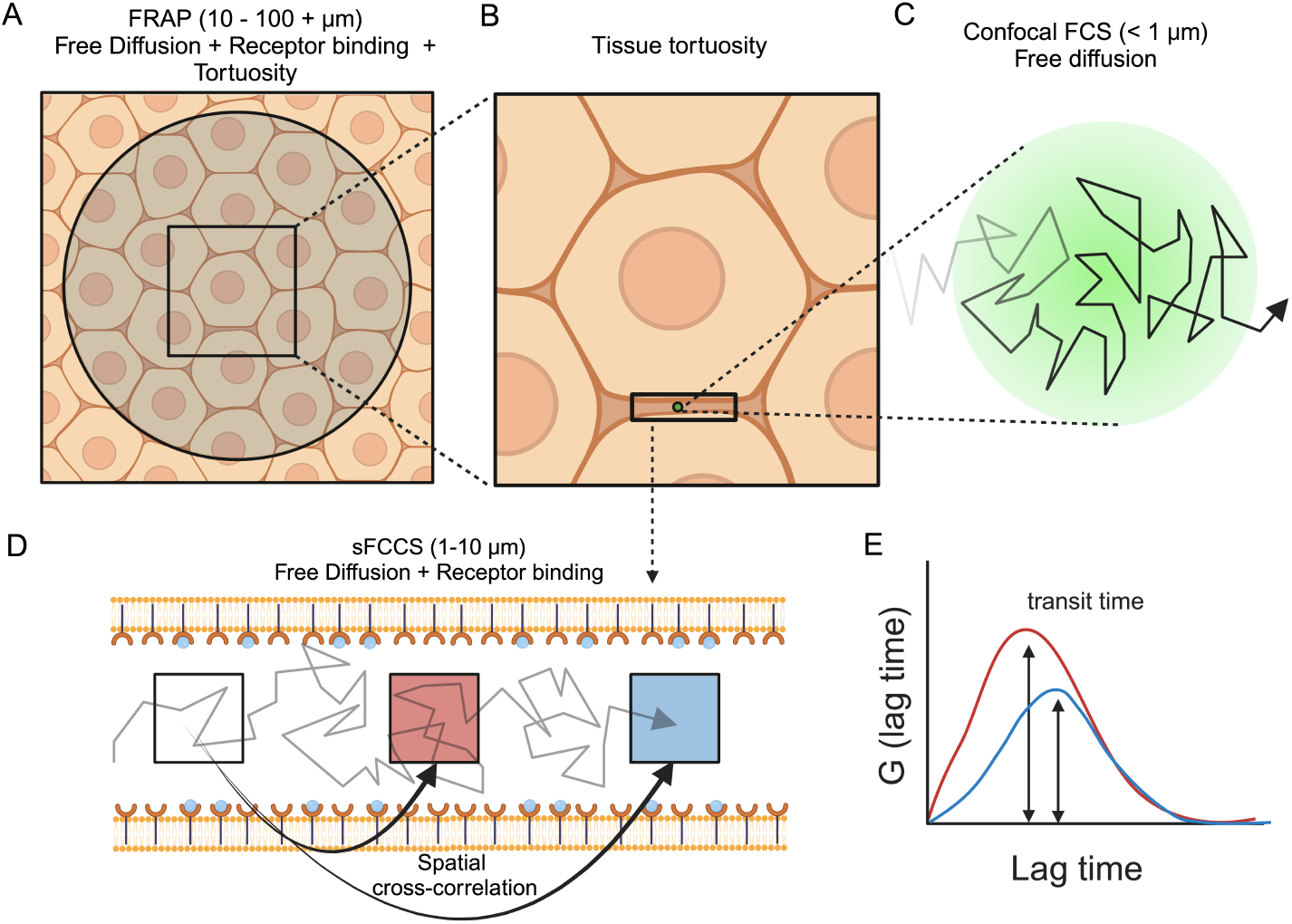
Hindered diffusion across different length scales: A.An illustration of the tissue space at the scale of 100 *μm*. FRAP measurements (area covered by the circle) are performed at this length scale, and they capture the combined influences of tortuosity and receptor binding to hinder the diffusion of morphogens. B.An illustration of the zoomed-in space of the tissue (square in A) capturing a single cell and its surrounding environment at the length scale of 20-30 *μm*. Morphogens diffuse and interact with the cell surface receptors in the interstitial spaces between cell membranes. C.An illustration of the zoomed-in space of the interstitial space (circle in B) capturing information from a diffraction-limited point in space (*<* 1 *μm*), as dictated by the observation volume of a confocal microscope. Morphogens may diffuse freely or interact with the ECM at this length scale, which can hinder their mobility at the local scale. D.An illustration of the interstitial space zoomed-in (rectangle in B), capturing the inter-membrane space between two cells at a scale of 10 *μm*. Morphogens can bind to the receptors on the membrane surface and transiently halt, hindering their diffusion across the mesoscale. E.An illustration of the application of the spatial cross-correlation approach to capture information about morphogen transits across two specific points within the inter-membrane space from Fig. 1 D. The red and blue ccf curves correspond to the respective transits of molecules measured from white to red and white to blue boxes respectively. The correlation peak captures the mean transit time of morphogens diffusing between a pair of points in space. The peak shifts to longer times with increasing separation distances since molecules take longer times to diffuse farther in space. Additionally, there is a reduction in correlation amplitude due to the reduced probability of finding the same set of molecules starting out from the initial point being considered for the spatial cross-correlation.

In this work, we address this information gap by probing the mobility transition from local to global length scales (0-10 *μm*) using spatial cross-correlations in Imaging FCS. Imaging FCS is an FCS modality which illuminates a thin cross-section of a sample and through fast camera readout, records intensity fluctuations caused by molecular movement for each pixel. By calculating and evaluating the pixel autocorrelation functions, it provides spatially resolved maps of diffusion coefficients [26]. Imaging spatial Fluorescence Cross-Correlation Spectroscopy (Imaging sFCCS) can be implemented with the same method but instead of auto-correlating the signal of each pixel, one cross-correlates pixels at different distances. Imaging sFCCS thus captures the transit time of molecules diffusing or flowing between any two points within the image (Fig. 1D, E). sFCCS has previously been established either by two focus FCS [27], [28], [29] or scanning FCS [30], [31], [32], [33] to measure transit times between two distinct points in space and thus obtain the diffusion coefficient or flow velocity of the molecule of interest. However, the information on spatial dynamics obtained from such techniques is limited to either two specific points or across a certain line in sample space. sFCCS performed on imaging-based modalities like pair correlation function (pCF) analysis [34], [35], [36], [37] and Imaging FCS [38], [39] allow for a wider spatial range for capturing dynamics between any set of pixels and in all directions. It thus helps obtain information about the cumulative influence of molecular interactions and barriers to diffusion in different directions of space, allowing one to correlate molecular mobility with the underlying micro-structure of the environment. Coupled with camera-based FCS [40], it has been used to probe space-dependent heterogeneity in diffusion dynamics *in vivo* [41], [42], [43].

The application of sFCCS on light sheet imaging data has already been used in the past to probe direction-dependent diffusion and hindrances on 2D cell membranes [44]. Here, we demonstrate the application of sFCCS in 3D within a single-plane illumination microscope (SPIM-sFCCS) to probe effective diffusion coefficients (*D*_*eff*_) over space in 3D environments. We modify the current 3D FCS fitting models of SPIM-sFCCS to account for signal crosstalk between pixels in a light sheet microscope for accurate *D*_*eff*_ estimation over space.

We apply SPIM-sFCCS to study the dynamics of the Nodal morphogen Squint (Sqt) to capture its scale-dependent *D*_*eff*_. Nodal signalling is essential for mesendoderm induction and left-right patterning in early zebrafish embryos [45], [46]. The dynamics and concentration gradients of Nodals have been extensively characterized due to their important role during gastrulation. Sqt is a morphogen since it can induce the expression of different genes across space in a concentration-dependent manner [47]. Sqt exhibits a *D*_*loc*_ of 40 - 60 *μm*^2^/s [16], [48] as measured by confocal FCS, and *D*_*glob*_ drops to as low 2 *μm*^2^/s when measured using FRAP. [10], [15].

While Sqt may not bind to the extracellular proteoglycans due to its residues giving it added stability and a long decay rate [10], [49] it does bind to cell surface receptors Acvr2b, 1a/1b, and the co-receptor OEP. It was shown that overexpression of OEP further slows down Sqt’s *D*_*loc*_ [21] and *D*_*glob*_ [12] and knockdown of a combination of receptors (Acvr 1b-a/b), increases Sqt’s *D*_*glob*_ [12].

Nodal signalling is active across 3-6 hours post fertilization (hpf) [50], [51] in a time window where the blastula tissue has large cells and when several tissue-level transitions occur that create different tissue densities [52]. Unlike in denser tissues seen at later stages, the blastula tissue, with visibly wide and varying inter-cell spaces available for measurement, makes it an amenable model environment for studying the mechanism of morphogen slowdown using SPIM-sFCCS.

Here, we first use diffusion simulations in channel-like spaces to understand and establish how the extent of cell-cell spacing and receptor binding can influence the scale and the value of *D*_*eff*_. Finally, we validate the simulation results experimentally. We measure *D*_*eff*_ of Sqt diffusing in the interstitial spaces between cells across different lengths and show that Sqt slows down and reaches a steady state diffusion coefficient value only at a distance greater than the intercell spacing. Moreover, knocking down the receptor Acvr2b or overexpressing the inhibitor Lefty2, increases the steady state *D*_*eff*_ of Sqt to a value similar to secreted EGFP, an inert non-interacting protein, implying that receptor binding hindered Sqt diffusion.

## 2 Results

### 2.1 SPIM-sFCCS

First, before applying SPIM-sFCCS in tissues, we derived a new theoretical fit model for data analysis. This became necessary as the crosstalk between the contiguous camera pixels leads to pseudo-autocorrelations that need to be taken into account for correct data evaluation (Fig. 2A, B).

**Fig. 2.**
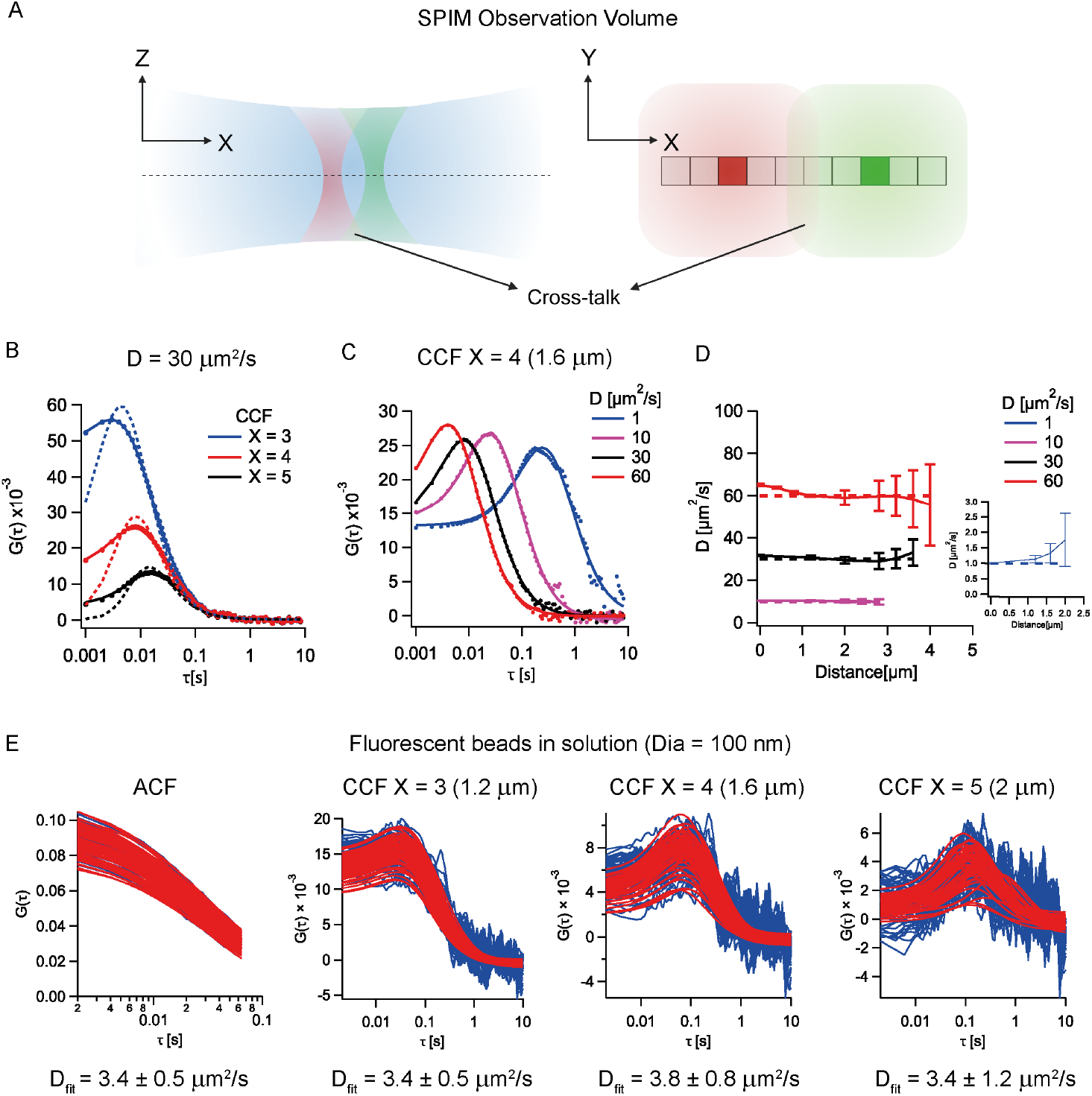
Addressing the issue of signal crosstalk across space in SPIM-sFCCS: A.Illustration of SPIM’s observation volume and how the overlap of the diverging sections of the observation volume in the axial direction causes an overlap in the detection volumes of a pair of pixels across space on the camera’s detector, resulting in signal crosstalk. Created in biorender.com B.Comparison of the modified fitting model and the previous model (dashed lines) for simulated data. The modified fitting model is able to capture the crosstalk of signal between pixels at short separation distances, unlike the previous model. C.Comparison of the average cross-correlation function from multiple correlations across space and the corresponding fits using the modified fitting model for different simulated diffusion coefficients for a certain separation distance. The modified model is able to capture the extent of the signal crosstalk component in the cross-correlation function for a range of diffusion coefficients tested. D.Fit data of the diffusion coefficient obtained from the average cross-correlations over space in simulated 3D free diffusion data with different diffusion coefficients. Error bars: SD of three independent simulations. The modified fitting model can capture accurate diffusion coefficients across a certain distance for a range of different diffusion coefficients. E.Comparison of the cross-correlation functions (blue) obtained from correlating multiple points in space and their corresponding fits (red) for different separation distances from freely diffusing fluorescent beads in water. The modified fitting model implemented on the GPU can capture the crosstalk of signal between pixels at short separation distances and provide constant diffusion coefficient values across space.

It is known that in multi-foci confocal FCS, overlapping observation volumes result in signal crosstalk between detectors [53]. The crosstalk, in this case, stems from the laser beam which propagates through the whole sample in the axial direction and produces out-of-focus fluorescence that is detected by neighbouring observation volumes [27], [54]. When cross-correlating neighbouring pixels, a pseudo-autocorrelation is seen as the pixels detect the same molecules at the same time. By enforcing a minimum distance between observation volumes, this crosstalk can be minimized.

In SPIM-sFCCS this problem is less severe as the light sheet has a finite thickness and thus less out-of-focus fluorescence is produced. Thus, the minimum distance between pixels that show no pseudo-autocorrelations is smaller. Nevertheless, as we use contiguous pixels in SPIM-sFCCS, the crosstalk between pixels can span over considerable distances and lead to a pseudo-autocorrelation component in the correlation functions (Fig. 2A). In our SPIM system, this signal crosstalk can be detected over a pair of pixels spaced almost 6 pixels (2.4 *μm*) apart (Supplementary Fig. 1A).

The extent of crosstalk increases with an increase in the span of PSF_*xy*_ due to a larger overlap in observation volumes (Supplementary Fig. 1B). The extent of crosstalk also increases with the thickness of the light sheet due to the increase in out-of-focus contribution from thicker light sheets (Supplementary Fig. 1C). However, the effects of signal crosstalk are not significant at larger separation distances (Supplementary Fig. 1D).

The fit model developed by Wohland *et al*. (2010)[40] did not account for the excess crosstalk, and the change in the PSF_*xy*_ along the z-direction. While it is adequate for fitting autocorrelations, it cannot fit the CCF functions.

We, therefore, developed a modified SPIM-FCS fit function based on Dertinger *et al*. (2007)[29] that accounts for this crosstalk by incorporating PSF_*xy*_ as a function of z, similar to the model for the confocal’s observation volume in Rigler *et al*. (1993)[55]. Since no analytical solutions could be found, the fit model was numerically integrated along the z-axis (Appendix A). The step size used in the numerical model was optimized to accurately fit ACFs and CCFs with D *>* 0.5 *μm*^2^/s (Appendix A Fig. A2). As fitting with numerical models is computationally expensive, we implemented data fitting on GPU, improving evaluation time 60-fold over CPU-based fitting.

The fit model was tested on 3D simulations for different distances and different D values. The model was able to capture the additional crosstalk component for different distances (Fig. 2B) and different diffusion coefficients (Fig. 2C). The fit model also gave accurate results within 20% of the set value for auto/cross-correlation data (Fig. 2D), with the range for accurate fits depending on the D value being measured (discussed in optimizing SPIM-sFCCS for *in vivo* measurements section).

Applying the model to fit spatial cross-correlations from SPIM-sFCCS measurements of fluorescent beads in solution yielded constant D across space, within 20% of the theoretically expected value while accounting for the increased pseudoautocorrelation component due to crosstalk (Fig. 2E). These results confirmed the model’s ability to accurately capture *D*_*eff*_ across 3D space.

### 2.2 Receptor binding simulations *in silico*

To effectively employ SPIM-sFCCS to probe the cause of morphogen slowdown across space, we first identified the physicochemical factors that influence the deceleration of morphogen diffusion. To that end, we simulated the mechanism of morphogen slowdown incorporating reversible receptor binding. We modelled the spaces between cells as channels of a certain width with the channel running along the x-axis. All simulated particles started across the middle of the channel at position (*x, y*) = (0,0). The particles spread with time, and their concentration along the x-axis was fit with a Gaussian function at regular intervals to capture *D*_*eff*_ from the resulting binding interactions (Illustration in Fig. 3A).

**Fig. 3.**
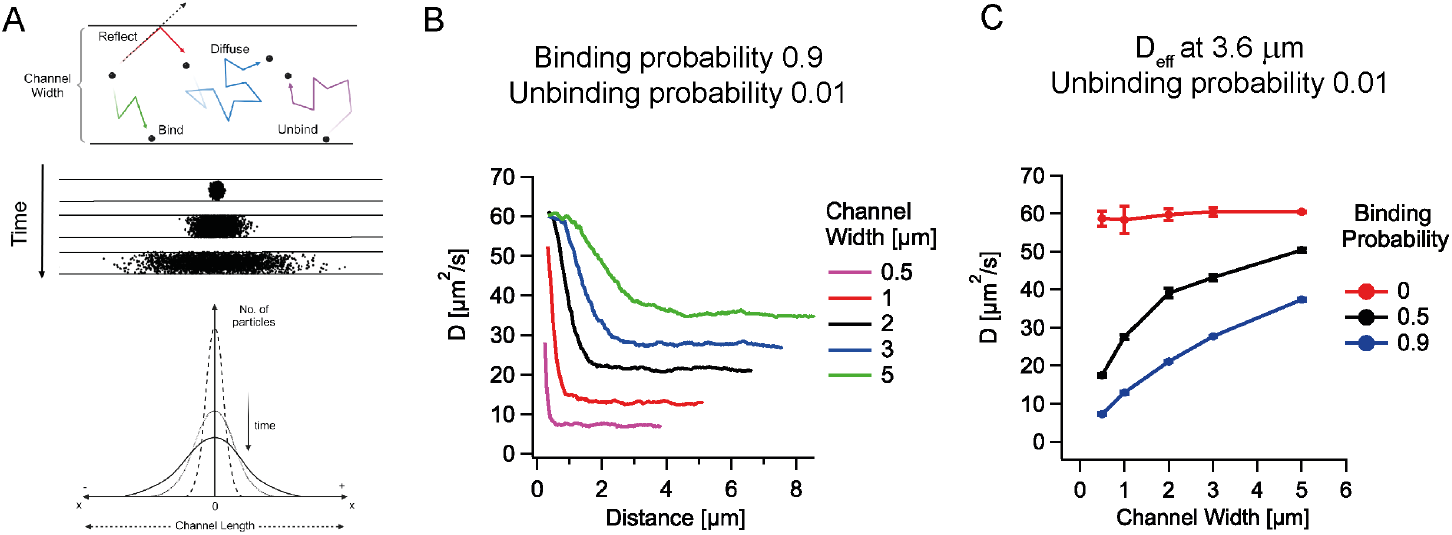
Simulations of binding-hindered diffusion in the inter-membrane spaces between cells: A.Illustration of how particles diffuse and bind to the walls of a channel in simulations. The effective diffusion coefficient over a certain period is calculated based on the distribution of particles across the length of the channel. The upper and lower illustrations were created in biorender.com B.Plot of the transition of effective diffusion coefficients over distance for different channel widths. The effective diffusion coefficient drops from the local value and reaches the steady-state value only after a certain distance dictated by the channel width. C.Plot of the effective diffusion coefficients obtained from different simulated channel widths and binding probabilities at a fixed distance of 3.6 *μm* (to compare with the results of Fig. 5B, C). Error bars are the SD of values from 3 independent simulation seeds. If the ligands experience a binding-based hindrance, the effective diffusion coefficient obtained at a steady state will be channel-width dependent.

Simulations showed a channel width-dependent diffusion coefficient, with smaller channels leading to a decrease in D (5 *μm*: 1.5 fold, 0.5 *μm*: 8 fold; see Fig. 3B). In all cases, *D*_*eff*_ converged to a steady state relatively quickly (0.01s-0.2s), with particles in narrow channels reaching a steady state quicker than in wider channels (Supplementary Fig. 2A). In the spatial domain, *D*_*eff*_ converged to a steady state value only at a distance greater than the width of the channel (Fig. 3B).

Increasing the binding probabilities progressively keeping the unbinding probability constant, reduced *D*_*eff*_ at steady-state (Supplementary Fig. 2B, D). For the 1 *μm* width channel, the highest binding probability set at 0.9 caused a slowdown of almost 5 fold. For the case of 0 binding probability, that is, when there was no binding, *D*_*eff*_ remained unchanged. Conversely, reducing unbinding probabilities, reflecting increased ligand-receptor affinity, significantly decreased the D (Supplementary Fig. 2C, E), with the extent of slowdown reaching almost 50-fold for the combination of 0.001 unbinding probability, 0.9 binding probability and 1 *μm* channel width. Zero unbinding probability did not allow for particle dispersal over space, emphasizing the necessity of reversible binding for ligand transport.

Overall, these simulations demonstrated that the combined influence of the binding parameters and the confined channel-like space was sufficient to slow down *D*_*eff*_ across space. Based on these simulations, measuring *D*_*eff*_ across a certain distance in a channel should yield a channel width dependent *D*_*eff*_ value in the presence of receptor binding. Reducing the binding levels should increase *D*_*eff*_, and in the case of no binding, *D*_*eff*_ should remain unchanged irrespective of channel widths (Fig. 3C).

### 2.3 Optimizing SPIM-sFCCS for *in vivo* measurements

Before applying SPIM-sFCCS in vivo, we developed data evaluation strategies to optimize the signal-to-noise ratio (SNR). We achieve this through spatial and temporal binning of pixels post-acquisition, where the binning range is determined by the particular range of concentrations and diffusion coefficients to be measured, and by the limited measurement time available due to the tissue dynamics of the embryo. In FCS, the SNR increases with the number of molecular transits recorded [56], [57]. Therefore, increasing measurement time reduces the % error of fit parameters and allows for accurate CCF fits over longer distances (Supplementary Fig. 3A, C). Similarly, increasing diffusion coefficients allows for more particle transits to be captured which also improves the signal for correlations (Supplementary Fig. 3B) and increases the fitting range (Fig. 2D, Supplementary Fig. 3D). This observation is advantageous for the current work since zebrafish ligand diffusion at the local scale ranges from 10-100 *μm*^2^/s [13]. However, for high D (60 *μm*^2^/s in Fig. 2D), estimations over short distances (β0-3 pixels) have poor accuracy due to the limited temporal resolution of the camera (β 1 ms).

Owing to the dynamic nature of the zebrafish blastula between 3-5 hpf, measurements with stationary interstitial spaces and constant inter-membrane spacing are limited in duration (30-120 s at 4 hpf, *<* 20 s at 5 hpf). These measurement times are insufficient to provide adequate transit statistics for SPIM-sFCCS measurements.

To improve the signal statistics for SPIM-sFCCS post-acquisition, we binned pixels around the two spatial regions of interest. Pixel binning increases the number of molecular transits being captured between the two points (Fig. 4A, Supplementary Fig. 3F). This, in turn, improves the quality of the cross-correlation functions and the accuracy and precision of the measured Ds (Fig. 4C and Supplementary Fig. 3E).

**Fig. 4.**
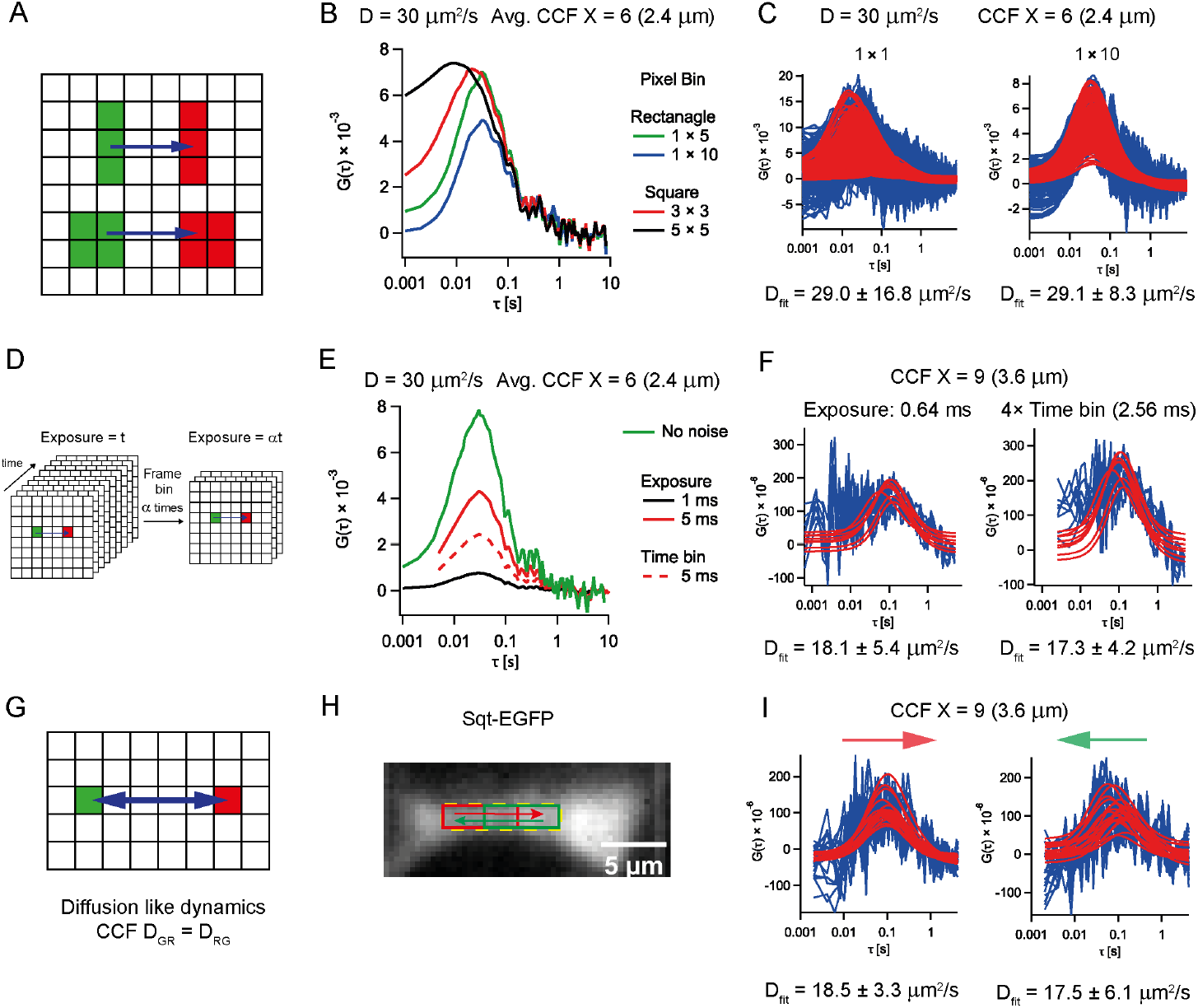
Signal improvement strategies for implementing SPIM-sFCCS *in vivo*: A.Illustration of binning pixels in space for SPIM-sFCCS. Camera-based Imaging-FCS allows for binning signals from multiple pixels in different ways to perform SPIM-sFCCS. B.Comparison between the CCFs calculated with rectangular and square binning of pixels from simulated data. Square binning causes the detection volume of pixels to overlap across the direction of measurement, causing the cross-correlations functions to have higher crosstalk than rectangular bins. C.Comparison of the raw cross-correlation function (blue) obtained and their corresponding fits (red) calculated with different pixel bin sizes from simulated data. With increasing binning size, the quality of correlations improves, resulting in better fits and low SDs in the obtained diffusion coefficient results. D.Illustration of binning frames across time for SPIM-sFCCS. Since a time series stack of image frames is collected for each measurement, consecutive frames can be binned (summed up) as an alternate way to increase exposure post-acquisition. E.Comparison of the average cross-correlation curve’s amplitude across different simulated signal levels. The 1 and 5 ms exposure data were from simulated stacks with 100,000 frames and 20,000 frames respectively and *N*_*noise*_ = 3. The highest amplitude is seen in the absence of added noise. The amplitude increases with increased acquisition exposure. The dashed curve is the average crosscorrelation curve from binning frames in time. Frame time-bin increases the amplitude of the crosscorrelation, but the increase is not as significant as with increased acquisition exposure. F.Example of frame time-binning used to improve the amplitude of spatial cross-correlations obtained from *in-vivo* measurements (raw correlations: blue, fits: red). Left spatial CCFs are obtained directly from the acquired image stack, of an Sqt-EGFP measurement while the CCFs on the right are from the same image stack frame binned 4*×*. The binning helps increase the amplitude of the CCFs to properly distinguish the CCF peak against the correlation amplitudes of background noise without affecting the CCFs’ peak positions and the fit diffusion coefficients significantly. G.Illustration of comparing the fit diffusion coefficient values in the forward and reverse directions. Equal forward and backward component values are required to ensure that the dynamics captured are diffusion-like and free from drift, flow, or other active dynamic components. H.An example of an ROI (region of interest) from an embryo expressing Sqt-EGFP to illustrate forward-backwards equivalence of measured effective diffusion coefficients. I.Spatial cross-correlation data obtained in the forward and reverse directions (raw correlations: blue, fits: red) from the data shown in H. The similar values indicate that the measured diffusion coefficients capture the true diffusive component of the dynamics without the influence of cell movement-based artifacts.

The amplitude of the correlation function decreases with increasing noise levels (Supplementary Fig. 3G) and increases with higher counts per molecule (Supplementary Fig. 3H) and higher exposure times (Fig. 4E). To improve SNR post-acquisition, we increased the acquisition exposure by binning frames in time (Fig. 4D). The correlation amplitude increases, although the improvement is not to the same extent as when acquiring with an equivalent high exposure value (Fig. 4E). Nevertheless, this approach not only allows the use of short and long exposures from a single measurement but also reduces the bias from fitting noise at small lag times for long-distance measurements on experimental data (Fig. 4F).

For *in vivo* measurements, to capture high-speed dynamics, we acquired signals with the fastest frame rate available for the measurement depending on the size of the ROI selected (0.5 ms-2 ms) and with higher laser power (60-600 W/cm^2^ compared to imaging (30-60 W/cm^2^) to capture sufficient counts per molecule. For long-distance measurements (3.6-5.2 *μm*), where cross-correlation amplitudes were low, we binned frames in time to obtain image stacks of 2-6 ms exposure time to increase SNR for improved data fitting.

Cellular movements can create convective flows that lead to deviations from simple diffusion (Supplementary Fig. 4A-right). As convective flows are directional, the CCFs will depend on the direction they are calculated for. Only if the forward and backward CCFs provide the same diffusion coefficient can one assume the absence of flow or directed transport (Fig. 4G). Discrepancies between forward and backward Ds indicated deviations from simple diffusion (Supplementary Fig. 4E, F, G) and such data were excluded.

Similarly, in the presence of barriers around the interstitial spaces, molecules diffusing into the barriers get reflected (Supplementary Fig. 4A-left). These reflected molecules contribute twice to the transit statistics, resulting in a wider CCF measured in the direction of the barrier compared to the one away from the barrier (Supplementary Fig. 4B, C, D). The approach of checking the equivalence of forward and backward Ds helps eliminate such effects.

In cases where the channel drifts along the z-axis, the forward and backward D may appear equal. However, the diffusion coefficient values are underestimated since the molecules take longer to transit across two points. Moreover, while we ensured that channel spacing remained constant within the analysis time window, we recognize that subtle changes in channel widths or the influence of the invisible axial spaces can lead to inaccurate estimations even if the bidirectional Ds match.

We found that at least 30% of all measured data had significant tissue movements, resulting in CCFs that could not be evaluated. 40% of measurements showed differences in the forward and backward diffusion coefficient, indicating convectional flow due to tissue movement. The remaining 30% of the measurements were evaluated.

We demonstrate the approach of comparing the forward and backward D of Sqt-EGFP along the length of the interstitial space between two cells that results in similar bidirectional D values when measured over a distance (Fig. 4H, I).

### *2*.*4 In vivo* results

The results from the simulations of particle diffusion in channel-like spaces with receptor binding suggest that the slowdown and the resultant steady-state *D*_*eff*_ are channel width-dependent, with the transition happening at a length comparable to the inter-membrane spacing between two cells. Having modified the fitting model and accounted for low signal and cell movements, we sought to verify the simulation results’ applicability to morphogen dynamics *in vivo*.

We initially compared *D*_*eff*_ of Sqt over a fixed distance of 9 pixels measuring 3.6 *μm* for channel-like spaces with different inter-membrane spacings. In Fig. 5A, B, it can be seen that for narrow spacings, *D*_*eff*_ are low, with the lowest spacing giving 10 β *μm*^2^/s. With increasing channel widths, *D*_*eff*_ measured increased till for spacings above 4 *μm* where *D*_*eff*_ values remained high with an average of 51 *μm*^2^/s. Furthermore, to compare the effect of the channel width and the extent of slowdown experienced in narrow spaces, we grouped the results from narrow spaces and wide spaces. We found that there was almost a 3-fold drop in *D*_*eff*_ for narrow channels (*<* 2.5 *μm*: β18 *μm*^2^/s) compared to channels of inter-cell spacings *>* 4 *μm* (β51 *μm*^2^/s). We also used these groups to measure the transition from *D*_*loc*_ to *D*_*glob*_ across the length of the channel (Fig. 5D). Following the receptor binding simulation results, in narrow channels (*<* 2.5 *μm*), *D*_*eff*_ dropped at 2 *μm* and remained relatively constant across space. For channel widths, greater than 4 *μm, D*_*eff*_ remained constant, with a marginal drop at 5.2 *μm* for a few channels. The channels with intermediate width showed a gradual slowdown till 3.6 *μm* and remained constant at 5.2 *μm* with *D*_*eff*_ higher than those of the narrow channels. Overall, this result recapitulates the simulation results to show that the scale of the slowdown depends on the inter-cell spacing.

**Fig. 5.**
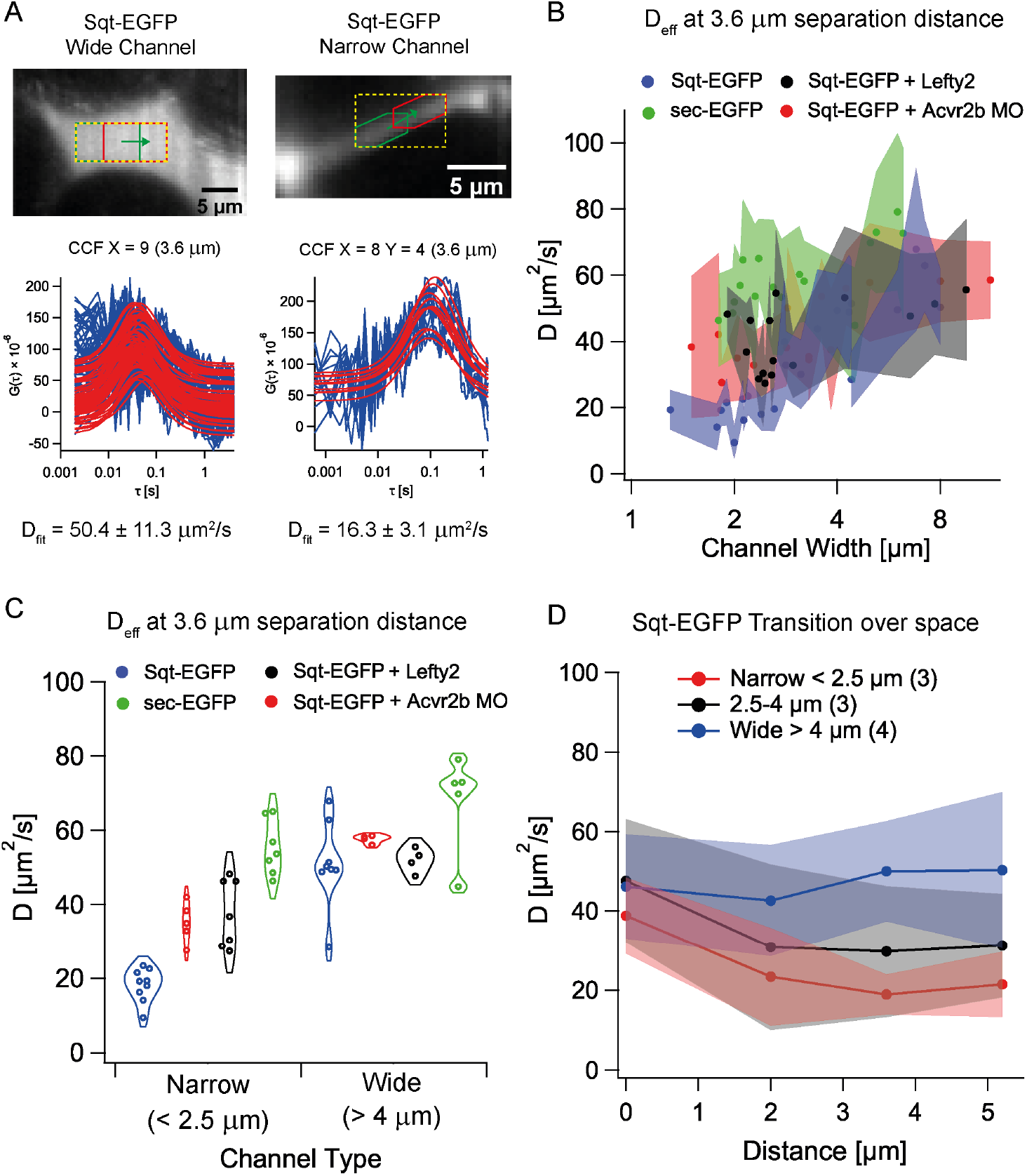
sFCCF measurements in channel like spaces *in vivo*: A.Comparison between the effective diffusion coefficients measured over 3.6 *μm* in a wide and a narrow channel-like space in Sqt-EGFP expressing embryos (raw correlations: blue, fits: red). Wider channel-like spaces allow for more space for molecules to diffuse over resulting in a faster effective diffusion coefficient while narrow spaces increase the interaction of the ligand with the cell surface receptors causing them to transiently bind while diffusing resulting in hindered diffusion across space. B.A comparison of the effective diffusion coefficients measured at a separation distance of ∼3.6 *μm* between sec-EGFP (*N*_*ch*_ = 16, *N*_*emb*_ = 9), Sqt-EGFP (*N*_*ch*_ = 22, *N*_*emb*_ = 15) and Sqt-EGFP in the presence of Lefty2 (*N*_*ch*_ = 15, *N*_*emb*_ = 8) and a morpholino against the Activin receptor 2b (Acvr2b) (*N*_*ch*_ = 15, *N*_*emb*_ = 9) from channels of different sizes (widths). Error range is the SD from multiple CCF fits from a specific channel. Note the effective diffusion coefficient of sec-EGFP for channels of small sizes (*<* 2.5 *μm*) is lower compared to that in wide spaces (*>* 4 *μm*). Sqt-EGFP has a much lower effective diffusion coefficient overall and exhibits a channel width-dependent diffusion coefficient, being slow in tight spaces and faster in wide spaces. The presence of Acvr2b morpholino and Lefty2 speeds up the diffusion of Sqt-EGFP in tight spaces while the effective diffusion coefficient in wide spaces remains similar to the non-treated case. C. comparison of the effective diffusion coefficients in A by grouping the data into narrow (*<* 2.5 *μm*) and wide (*>* 4 *μm*) channel sizes (widths) based on the separation distance of *β* 3.6 *μm*. The morpholino against the Acvr 2b receptor and Lefty2 speed up the slow Sqt-EGFP in narrow channels, while its effect is not significant in wide spaces. Sec-EGFP has a higher diffusion coefficient in narrow and wide spaces compared to Sqt-EGFP in morpholino-treated and untreated cases but still exhibits a relative slow-down in narrow spaces compared to wide spaces. D.Measurement of the transition of diffusion coefficients for Sqt-EGFP across space for different channel sizes (widths) (*N*_*ch*_ = 10, *N*_*emb*_ = 9). The average D value and SD from the pooled channels were taken and plotted (channel number for each group in bracket) with the error range being the SD of the pooled results. In tight spaces, within the measured distance, the effective diffusion coefficients are high locally and slow down when measured at a distance larger than the channel size (width) and remain mostly constant. In contrast, the diffusion coefficient in wide spaces (*>* 4 *μm*) is high and constant across space within a margin of error.

Since Sqt is known to bind to Activin receptors, we investigated whether modulating receptor binding could influence *D*_*eff*_ over space. For this, we knocked down Acvr2b, a receptor known to bind to Sqt *in vivo* [48] using a morpholino [58] and measured *D*_*eff*_ at 3.6 *μm* to compare with the scatter plot of Sqt-EGFP’s diffusion. We found that *D*_*eff*_ in narrow spaces increased compared to native Sqt-EGFP values, with a wide distribution of observed *D*_*eff*_ values between 30-60 *μm*^2^/s (Fig. 5B). Grouping the narrow spaces together to compare with Sqt-EGFP showed a significant overall increase in the diffusion coefficient by about 2 fold (β18 *μm*^2^/s to β35*μm*^2^/s), while the wide space *D*_*eff*_ remained relatively unchanged compared to Sqt-EGFP (β51 to β57 *μm*^2^/s)(Fig. 5B).

Leftys are known inhibitors of Nodal morphogens. They act by either directly binding to the Nodal ligand or by competitively binding to receptors to prevent Nodal signalling [49], [59], [60]. In zebrafish, Leftys preferentially inhibit Squint [61] and exhibit higher diffusion coefficients compared to Squint and Cyclops across tissue spaces [10]. We previously demonstrated that Squint binds to Leftys in the extracellular space [48]. Here we examined whether overexpressing Leftys could speed up Squint’s diffusion since both direct interaction of Squint and Lefty and Leftys blocking available receptor binding sites should overall reduce the transient binding of Squint. Accordingly, we measured Sqt-EGFPs *D*_*eff*_ in the presence of Lefty by overexpressing Lefty2-mCherry in the extracellular space. Similar to the Acvr2b receptor morpholino result, we found a two-fold increase in the diffusion coefficients for the Sqt-EGFP with Lefty2 overexpression in narrow spaces compared to native Sqt-EGFP values (β18 to β38 *μm*^2^/s) while the *D*_*eff*_ in wide spaces remained relatively unchanged (β51 to β52 *μm*^2^/s, Fig. 5B, C).

It should be noted that in the transition from wide to narrow channels, the dimensionality reduction from 3D (wide channels) to 2D (narrow channels) leads to a reduction of the mean squared displacement from 6Dt to 4Dt [62], [63]. As our fit model cannot take account of this transition, this results in an underprediction of D by 30%. We tested this prediction using sec-EGFP and obtained apparent reductions in D on that order of magnitude for narrow channel spaces (Fig. 5B, C). Since this effect is also applicable to the diffusion of Sqt, the 30 % underestimation of the diffusion coefficient in narrow spaces should result in a value of 35-40 *μm*^2^/s. However, the values measured for Sqt in narrow spaces are as low as 10 *μm*^2^/s and average around 18 *μm*^2^/s. A value of β35-38 *μm*^2^/s is recovered only upon Acvr2b knock-down or Lefty2 overexpression.

Overall, these results suggest that modulating binding affinity does indeed influence *D*_*eff*_ of morphogens across space, and therefore, these morphogens experience hindered diffusion through binding, acting at the length scale of the intercell spacing.

To further test the validity of our observations from Sqt-EGFP diffusion, we used SPIM-sFCCS to measure the *D*_*eff*_ of BMP2b-sfGFP across space. BMPs and Nodal signalling work synergistically to pattern the early zebrafish embryo [64]. Similar to Squint, BMP2b has also been shown to exhibit a difference in *D*_*loc*_ and *D*_*glob*_ values [11], [65], [66] and thus, can be useful to compare with Sqt results. From the measured *D*_*eff*_ values from a limited set of narrow and wide channels (Appendix B), we found a distinct difference between the channel types, with narrow channels significantly restricting ligand diffusion, slowing the molecules down by almost 4 fold compared to that in wider channels *in vivo* (56 to 15 *μm*^2^*/s*).

## 3 Discussion

Quantifying *in vivo* morphogen dynamics is crucial for understanding morphogenesis. However, current techniques, including FCS, SPT, and FRAP, probe different non-overlapping spatial scales and thus capture the spatial scale-dependent dynamics incompletely. To capture the full gamut of effects of tortuosity, extracellular matrix and receptor interactions over all length scales, therefore, requires techniques that can bridge the transition from local to global dynamics.

Here, we employ SPIM-sFCCS with improved fit and analysis procedures to measure *D*_*eff*_ *in vivo* across intermediate-length scales. Due to the large detection volume per pixel and the extent of the light sheets, crosstalk between neighbouring pixels is strong and extends over a wide range. This was previously recognized, and solutions for deconvolution were developed [67]. However, for SPIM-sFCCS, this leads to enhanced crosstalk that cannot be accounted for by previously derived analytical models [40]. Correct D values can be extracted from these models but at the cost of an empirical calibration, which will overestimate the observation volume. This approach, however, fails for SPIM-sFCCS, and the derivation of a new model became necessary.

The disadvantage of this newly derived model is that no analytical solution can be found, and numerical integration is required to describe the fit function, which slows down data evaluation. But it should be noted that at the same time, this makes the model versatile as any PSF model can be included in the numerical function, allowing the inclusion of even experimental PSFs [68]. An alternative approach to data fitting is the evaluation by convolutional neural networks [69]. Finally, we demonstrated that with the developed numerical fit model, we can extract correct Ds over distances ranging from at least 0-10 *μm*, spanning the accessible length scales between FCS and FRAP.

We used simulations to provide predictions of how *D*_*eff*_ changes over distance with receptor interactions. For this purpose, we used single effective binding and unbinding probabilities of a generic receptor. We recognize that several receptors and co-receptors of different time-varying concentrations are involved, and their individual effect on receptor binding-based hindered diffusion can be hard to predict. Nevertheless, their overall influence can be captured within these generic simulation parameters, and the predictions should at least qualitatively capture the effect of the receptors’ interaction on morphogen dynamics.

Using this simplified model, receptor interactions can be tuned by either increasing the ligand-receptor affinity (binding and unbinding probabilities) or by changing the receptor concentration. Assuming a constant surface concentration of receptors on the cell membranes, the latter is equivalent to changing the spacing between the cell membranes that form the channels through which the morphogens diffuse. These simulation results are consistent with those of Kuhn *et al*. (2022)[21] and Zhu *et al*. (2024)[14], who showed how binding affinity, receptor concentration, and tightness of space influenced global diffusion. Furthermore, we predicted that the transition from free to diffusion hindered by receptor interactions occurred at length scales on the order of the inter-membrane distance.

We then proceeded to apply SPIM-sFCCS in the living embryo. With the blastula tissue undergoing phase transformation at 4 hpf [52] causing global cell drifts and the Nodals’ ability to induce active cell movements [70] by activating the cytoskeleton polymerization [71], it was important to distinguish tissue movement artifacts, convection induced by tissue movement, and diffusion. In addition, the complexity of the 3D tissue topography and dynamics led to a wide distribution of D values in general. For the optimization of SPIM-sFCCS in a highly dynamic tissue, we needed to find a compromise between limited measurement time and achievable SNR. As tissue moves fast in the early embryo and can corrupt the correlation functions, we could not measure over long times (Mean measurement time ≈ 90 s). The resulting low SNR was mitigated by pixel binning and frame binning to obtain appropriate spatial cross-correlations and quantify *D*_*eff*_. These strategies were crucial for the long-range measurements of the 3.6-5.2 *μm* separation distances shown.

Taking into account these limitations, we applied SPIM-sFCCS to the measurement of the scale-dependent diffusion coefficient of the nodal morphogen Sqt, dependent on receptor concentration and channel width. Our measurements supported the simulation predictions, and we showed that even after taking into account the effects of dimensionality reduction, Sqt slows at least 3-fold over the length of tight channels while remaining unperturbed in wide spaces. Both, Acvr2b receptor knock-down and Lefty2 overexpression increased *D*_*eff*_ of Sqt in tight spaces while not affecting the diffusion in wide spaces, demonstrating the channel-width-dependent effect of receptor interactions. Furthermore, SPIM-sFCCS measurements on BMP-2b also showed differences in the *D*_*eff*_ measured in narrow and wide spaces, thus suggesting the general applicability of the mechanism of space-dependent binding-hindered diffusion for morphogens.

An inert control, secreted EGFP, did show an apparent β 30% slow down in D for very narrow channels due to dimensionality reduction of diffusion from 3D to 2D. It should be noted that this is not an actual decrease of D of EGFP but the inability of the fit model, which was derived for the 3D case, to account for the varying changes when the sample space transitions from a 3D to a 2D space. There was no channel size-dependent diffusion and no effect of receptor binding, unlike Sqt, which showed a much stronger reduction of D over space.

A more detailed analysis of the experimental D values of Sqt in channels of different widths showed a gradual decrease of D over distance, which depended on the channel width, as predicted by the simulations. To our knowledge, this is the first scale-dependent measurement of *D*_*eff*_ of a morphogen in a living embryo, and it demonstrates the power of SPIM-sFCCS.

These *D*_*eff*_ measurement results are consistent with previously reported receptor binding hindered diffusion of ligand measurements *in vivo* [12], [14], [18], [19], [21]. Our group also previously demonstrated receptor binding hindered diffusion in tight interstitial spaces of the 24-48 hpf zebrafish cerebellum for Wnt3, where knocking down co-receptor LRP-5 increased *D*_*glob*_ values [18]. Furthermore, it was shown that BMP4 gradients in mice formed due to an interplay of geometry-constricted ligand diffusion in narrow interstitial spaces and restricted basolateral localization of receptors [72].

Our results also lead to further predictions. Endogenous nodal signalling is preferentially active at the dorsal margin of the blastula [73]. When these margins still have intact cell-cell adhesions [74], the inter-membrane space in such a region might be much below 100 nm and, therefore, below the diffraction limit. Such tight spaces should cause a significantly greater slowdown for ligands. Due to the diffraction limit, we can’t quantify the tight spaces. Nevertheless, based on our results, we extrapolate that *D*_*eff*_ values in these denser tissue regions will be much lower than our results for 2 *μm* channels of 10-20 *μm*^2^/s.

Based on our scale-dependent *D*_*eff*_ results, we illustrate a plausible mechanism by which molecules slow down from local to global length scales in Fig. 6. The receptor binding-based slowdown happens within the interstitial space between a pair of cell membranes (Fig. 6A). Beyond the intermembrane spacing distance, the resultant *D*_*eff*_ value is maintained across the length of the interstitial space. After transversing the intermembrane space, the molecules experience the tortuosity of the tissue (Fig. 6A). They must circumvent the cell and may get confined in the void spaces between cells, which delays their dispersal and causes a further slowdown. We predict that the transition from a fast *D*_*loc*_ to a *D*_*eff*_ reduced by receptor binding, and if applicable by interactions with extracellular matrix components, occurs within one cell diameter itself. The final *D*_*glob*_ beyond the scale of a single cell, which helps shape the concentration gradient, is then determined by tissue tortuosity (Fig. 6A). Potentially, this can explain how Nodals’ and other morphogens’ mobility is regulated to form concentration gradients *in vivo*.

**Fig. 6.**
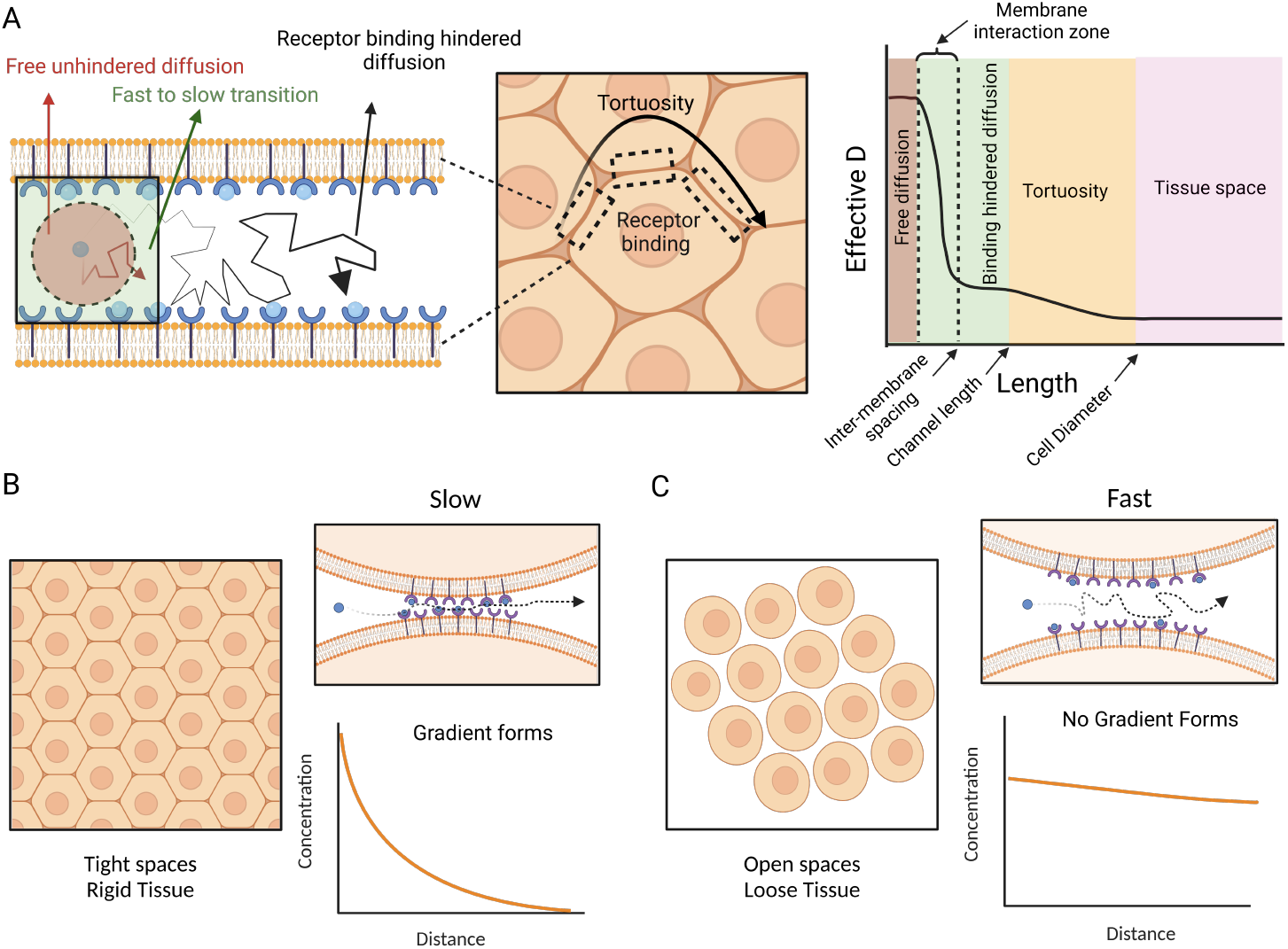
Plausible mechanism of morphogen slow down across space for formation of concentration gradients: A.Illustrations to show how interactions at different length scales influence the diffusion of morphogens. At the scale within the inter-membrane space, morphogens exhibit free diffusion or ECM interaction hindered diffusion. Beyond the length scale of the inter-membrane spacing and within the channel-like interstitial space, morphogens experience the influence of receptor binding. Beyond the intermembrane space, the tortuosity of the tissue requires the molecules to diffuse along a longer route around the cell’s perimeter, further slowing molecules down. The entire transition from a local fast to a global slow can thus take place within the scale of one cell diameter. B.The limited spaces between cells in a rigid tissue increase the propensity for morphogens to bind to their receptors transiently, causing them to slow down significantly and form gradients. C.As the embryo advances to the epiboly stage, the tissue undergoes a phase transition that causes the cells to become loosely packed and permits the rapid drainage of molecules. Gradients may no longer form or exist at this point. Figure created in biorender.com

With the tissue spaces in the blastula actively changing at the pre-gastrulation time window where nodal signalling is active (3-6 hpf), a question of how a morphogen gradient is maintained despite tissue level changes spaces arises. The timing of nodal expression is very precisely regulated between 3.5-5 hpf [50], [51], [73]. At a 4 hpf transition, the blastula tissue undergoes a phase transition, creating wide inter-cell spaces and allowing for faster diffusion [52]. However, the cell-cell adhesions in the margin spaces remain intact due to the persistence of non-canonical Wnt11 signalling [74]. We speculate that these dense margin spaces can sufficiently slow down Sqt’s diffusion to create and maintain a concentration gradient (Fig. 6B). When these cell-cell adhesions are lost at 5 hpf, the open spaces created will not allow for rapid deceleration of Squint and thus prevent gradient formation (Fig. 6C). Moreover, the spaces will also allow the pre-repressed Leftys [50] activated this time to diffuse quickly and inhibit further nodal signalling. Nonetheless, to test this mechanism, the changes in tissue packing of the margin spaces and the resulting ligand dynamics need to be correlated with the downstream response times of these ligand signals in future work.

The implementation of SPIM-sFCCS provides a useful tool for investigating the diffusion coefficients in live embryos in great detail. It achieves access to the intermediate range of up to 10 *μm*, which is not accessible by more traditional measurement techniques. Our data from both Sqt and BMP-2b indicates a slowdown of morphogen diffusion with distance within the extracellular space that is dependent on receptor concentration and affinity as well as intercell spacing, leading to an effective reduced *D*_*glob*_ already at a distance comparable to the diameter of a cell. We expect that with further technical developments, detailed maps of diffusion coefficients can be measured in live embryos, providing a much more detailed picture of morphogen transport than is available today.

## 4 Materials and Methods

### 4.1 Simulations for diffusion and binding in narrow channels

The diffusion of morphogens in a developing embryo is hindered not just by the geometry of the tissue but also by binding and unbinding to receptors on the cell walls. We simulated a simplified version of this situation in 2D by creating a channel of width *d*_*width*_ bounded by two parallel lines parallel to the x-axis representing cell borders. All particles started diffusing from the centre point of the channel. The simulation time step was set to ∆*t* = 0.01*ms*, and the diffusion coefficient was *D* = 60*μm*^2^*/s*. The time step of 0.01 ms ensures a short step size of 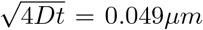 for 60 *μm*^2^/s so that particles will require multiple steps to cross the channel. Particles perform a random 18 walk within the channel with step size ∆*r* in *x* and *y* direction drawn from a Gaussian distribution (mean 0, variance 2*D*∆*t*). Once a particle hits one of the walls, it can either bind with probability *p*_*bind*_ or be reflected as determined by drawing a number from a uniform distribution between 0 and 1 and comparison to *p*_*bind*_. A bound particle has a probability *p*_*unbind*_ to unbind and continue diffusion in the next simulation step, determined by drawing a number from a uniform distribution between 0 and 1 and comparison to *p*_*unbind*_. As all particles start at the centre point of the channel, the distribution of the particles along the x-axis, that is, along the channel, provides a measure of the effective diffusion coefficient (*D*_*eff*_) resulting from the free diffusion and transient binding-unbinding dynamics over space. The resultant distribution of the spread of the particles along the length of the channel was fit every 100-time steps using a Gaussian fit function:

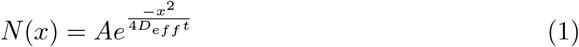

The number of particles in each simulation was scaled according to the width of the channel as 4000 particles/*μm* of channel width. This ensured that there were sufficient particles to create a Gaussian distribution that could be fitted appropriately. Furthermore, the length of each bin size for the histogram capturing the distribution of particles was set to 0.1 *μm* over the first 0.2 seconds and later to 0.5 *μm*. At the initial time points of the simulation, the particles do not cover a large distance. Therefore, the bin size needs to be small enough to capture minute differences in the particle distribution over space. At later times, when the particle distribution over space is sparse, coarser bins can capture the global spread.

The distance particles transversed along the x-axis within each time interval (1 ms) were calculated using the following equation.

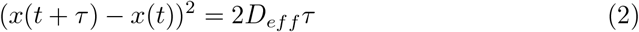

Here *x*(*t*) is the position at time t, and *τ* is the time interval for each measurement of *D*_*eff*_. All simulations of channels were performed in Mathematica version 12.3 (Wolfram Research, Illinois, USA).

### 4.2 SPIM-FCS

The light sheet microscope system and acquisition settings were based on the details presented in Krieger *et al*. (2015)[26] and are briefly described here. A 488 nm laser (Cobolt 06-MLD 488nm 0488-06-01-0100-100, Cobolt AB, Sweden), required to excite EGFP, was coaligned with a 561 nm laser (Cobolt 06-DPL 561nm 100 mW 0561-06-91-0100-100, Cobolt AB, Sweden), used to excite mApple/mCherry. The beams were directed into a single-mode optical fibre (kineFLEX-P-3-S-405..640-1.0-4.0-P2, Qioptiq, USA) which expanded the beams by 1.3 times. The output beams were passed through a beam expansion (*f*_1_ =30 mm AC254-030-A, *f*_2_ = 60 mm AC254-060-A; Thorlabs Inc., USA) to obtain a further 2× expansion, which was required to fill the back aperture of the illumination objective. The beams were then passed through an achromatic cylindrical lens of 75 mm focal length (ACY254-075-A; Thorlabs Inc., USA) followed by an illumination objective (SLMPLN 20 × /0.25; Olympus, Tokyo, Japan). The light sheet thickness obtained for the 488 nm beam had a 1*/e*^2^ radius of approximately 1.1 *μm* and was maintained for all SPIM-sFCCS measurements. The light sheet waist was aligned to coincide with the optical axis of the detection objective (LUMPLFLN 60× /1.0, Olympus, Tokyo, Japan). The detection objective was placed in a mounting hole on one side of a custom-made sample chamber and mounted on a piezo objective scanner (P-721 PIFOC; Physik Instruments, Germany) to control the objective with nanometer precision. The sample was mounted on a motorized stage with three linear positioning systems (Q-545 Q-Motion Precision Linear Stage; Physik Instruments, Germany) with piezo motors for the three axes and one rotation stage (DT-34 Miniature Rotation Stage; Physik Instruments, Germany). The emission obtained by the detection objective was passed through a bandpass filter (FF03-510/20-25, Semrock, USA) to capture EGFP emission and FCS, or a dual-band bandpass filter (FF01-512/630-25, Semrock, USA) to check for mApple/mCherry with EGFP co-expression, and was projected onto an EMCCD camera (Andor iXon3 860, 128 × 128 pixels; Andor Technology, Belfest, UK) by a tube lens (LU074700, f = 180 mm, Olympus, Tokyo, Japan). A flip mirror was used to alternatively project the image onto a sCMOS camera (OCRA-Flash4.0 V2; C11440; Hamamatsu, Shizuoka, Japan) with a large field of view to visualize the global position of the embryo and to decide on the region of interest.

Time series stacks were obtained in TIFF format from the EMCCD by the software Andor Solis for Imaging (version 4.31.3037.0), and were subjected to SPIM-sFCCS analysis depending on the acquisition parameters of exposure and measurement time used.

### 4.3 Free diffusion simulations for SPIM-sFCCS

Simulations of free diffusion in SPIM (3D) were performed in Imaging FCS 1.52 [75], an ImageJ (http://imagej.nih.gov) plugin building on the 2D TIRF simulations described in Sankaran *et al*. (2013)[76] using the simulation parameters given below. Briefly, we assumed that fluorescence signals were collected through a 60×, NA 1.0 microscope objective for SPIM. The emitted signal was imaged onto a camera with a pixel size of 24 *μm* (object space pixel size = 400 nm), and of which a 20× 20 pixel area is observed (corresponding to 8 × 8 *μm*^2^ in SPIM). To ensure freely fluctuating particle numbers in the observed area/volume, the simulation space was chosen to be 3 times larger in x and y dimensions than the observed area and additionally 20 times larger in z direction than the light sheet thickness (1/*e*^2^ radius) with the observation volume centred in the simulation space. A defined number of particles was distributed randomly in space. For every simulation step of time ∆*t*, particles were displaced in each of the three dimensions by a random step size drawn from a Gaussian distribution with mean 0 and variance 2*D*∆*t*, where D is the diffusion coefficient. If a particle left the simulation space, a new particle was created randomly on the border of the space. The time step ∆*t* was chosen to be 1*/*10^*th*^ of the exposure time T of the camera to account for particle movement during image acquisition. At each simulation step, each particle emitted a number of photons sampled from a Poisson distribution with a mean of *F* (*z*)*cps*∆*t*, where *cps* represents the molecular brightness in counts per particle per second and *F* (*z*) is a scaling factor taking into account the axial dependence of the excitation intensity. The emitted photons were distributed on the camera with a probability distribution as defined by the point spread function (PSF) of the microscope objective in the *xy* plane. The PSF was approximated as a Gaussian function with a 1*/e*^2^ radius of

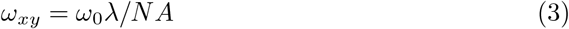

here *ω*_0_, a constant, was obtained from a calibration experiment as described in the experimental section. The scaling factor *F* (*z*) is given by a Gaussian distribution of the light sheet intensity in the Z direction.

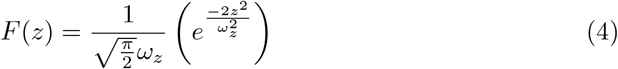

where *ω*_*z*_ is the 1/*e*^2^ value of the light sheet’s thickness. As particles outside the focal plane will result in a wider distribution of the photons on the camera, photons were spread over a PSF with

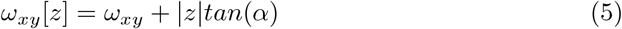

The detected counts were summed up and projected onto a 2D image every 10 simulation time steps (*T* = 10∆*t*), representing one frame in the image stack. A maximum of 500,000 frames were simulated for SPIM-sFCCS analysis. To simulate the effect of camera noise, spurious counts were randomly drawn from a Gaussian distribution (mean 0, standard deviation *σ* = *N*_*noise*_) and added to each pixel in each frame.

For simulation results shown in Fig. 2B, C, D; Fig. 4B, C, E and Supplementary Fig. 3, image stacks were simulated and analyzed with 5000 particles; 100,000 frames; exposure time = 1 ms; 30,000 cps; PSF_*xy*_ = 515 nm ; PSF_*z*_ /light sheet thickness = 1.13 *μm*; 1×1 binning unless mentioned otherwise for the specific sub-figure.

### 4.4 Fluorescent bead measurements procedure

Diffusion measurements of fluorescent beads in solution were performed based on the procedure described previously [26]. Briefly, fluorescent beads (FluoSpheres, Invitrogen) with a 100× nm diameter and emitting fluorescence signal at 515 nm wavelength were diluted 100× of the stock solution in water and used here for Imaging-FCS measurements. The bead solution was sonicated for 30 minutes prior to measurement. 50 *μl* of the beads solution was enclosed in heat-sealed FEP (fluorinated polyethenepropylene) bags with a film thickness of 10–25 *μm*. The sealed bag was mounted into the sample chamber, and the signal from the fluorescent beads was captured for SPIM-sFCCS analysis. 80,000 frames with an exposure of 2 ms for a laser intensity of 12 W/*cm*^2^ were acquired for the measurements.

### 4.5 mRNA preparation, injection, zebrafish mounting and SPIM imaging

The Sqt-EGFP, sec-EGFP and the Lefty2-mCherry plasmid constructs used here were previously published by Wang *et al*. (2016) [48]. The plasmids were linearized using the NotI restriction endonuclease (New England Biolabs, USA). mRNA was transcribed from the plasmid using a transcription kit (AM1340 mMESSAGE mMACHINE SP6 Transcription Kit, Thermofisher Scientific, USA) and purified using Dynabeads mRNA Purification Kit, (Thermofisher Scientific, USA). The morpholino against Acvr2b (GeneTools, USA) is based on the antisense sequence (GCAGAGAAGCGAACATATTCCTTT) in Albertson *et al*. (2005)[58].

Zebrafish embryos were obtained from the Comparative Medicine facility at the National University of Singapore. All experiments were performed following an approved protocol (IACUC #BR18-1023) by the Institutional Animal Care and Use Committee at the National University of Singapore.

10-15 pg of Squint-EGFP or sec-EGFP mRNA were injected into the single cell of the eight-cell stage embryo for SPIM-FCS measurements. This concentration is higher than 5-10 pg used for confocal FCS previously [48] to ensure that sufficient signal was captured through the SPIM system and the EMCCD camera. Higher concentrations may increase SNR for imaging but make it harder for the fluctuations in intensity with respect to the background intensity to be appropriately detected which is crucial for FCS.

For Morpholino injection, a cocktail of the Morpholino and PMT (plasma membrane targeting sequence)-mApple mRNA was injected in each of the cells of the two-cell stage to ensure that only fertilized embryos were injected. The concentration of the injected morpholino was 2.5 *μg*/*μl*. This is in the same order 4 *μg*/*μl* used to knock down the expression of Acvr2b in the report by Albertson *et al*. (2005) [58]. Along with it, 50 pg of PMT-mApple [77] was injected to visualise where the injected cocktail with morpholino acted in the embryo and ensure that the cell membranes flanking the regions measured had their Acvr2b receptors knocked down.

Lefty2-mCherry was similarly injected in each of the cells of the two-cell stage in the embryo resulting in a total injection load of β 60 pg to ensure adequate over-expression across the extracellular space.

The Bmp2b-sfGFP construct used here was previously published by Pomreinke *et al*. (2017)[11]. The plasmid was linearized using the NotI restriction endonuclease (New England Biolabs, USA). mRNA was transcribed from the plasmid using a transcription kit (AM1340 mMESSAGE mMACHINE SP6 Transcription Kit, Thermofisher Scientific, USA) and purified using the Zymoclean RNA Clean & Concentrator kit (Zymo Research, USA).

10-15 pg of Bmp2b-sfGFP was injected into the single cell of an eight-cell stage embryo for SPIM-FCS measurements.

The embryos were screened at 3 hpf for EGFP/sfGFP/mApple/mCherry expression in a fluorescence stereo microscope. For SPIM-FCS measurements, the selected embryos were dechorinated and were suspended in 1% low melting agarose (UltraPure™ Low Melting Point Agarose, 16520100, Thermofisher Scientific, USA) in a Fluorinated Ethylene Propylene (FEP) tube of 1.1 × 1.5 mm^2^ cross-section (FT 1.1× 1.5, Adtech Polymer Engineering, England). The embryos were mounted and imaged between 3.5-5 hpf before severe cell movements made FCS analysis difficult.

To minimize imaging artifacts, the embryo was oriented in a way that the blastula space, based on the region of interest for imaging, was oriented to be closest to one face, and the animal pole of the embryo was placed as close to the wall of the FEP tube. Essentially, to have the embryo’s animal pole positioned within one quadrant of the cylindrical tube. This ensured that the light sheet entering the tube could illuminate a plane of the embryo’s tissue optimally and that the detection objective could capture the fluorescence emission without any losses during the propagation through the tube and agarose.

Initially, the embryo was imaged on the sCMOS to visualise the embryo orientation and choose a broad region of interest (221.9 *μm ×* 221.9 *μm*). Once chosen, we switched to the EMCCD camera to image a smaller ROI (51.2 *μm ×* 51.2 *μm*). To find appropriate channels like inter-membrane spaces, the tissue space was surveyed with low laser intensities (30-60 W/cm^2^) and high exposures (50-100 ms) to reduce photobleaching. We avoided ROIs showing cell movements, shadows, or scattering artifacts. Once the ROI was chosen for SPIM-sFCCS and the field of view was cropped to allow for high frame rates, image stacks with about 100-150k frames were recorded with low exposures (0.4-2 ms) and high laser intensities (60-600 W/cm^2^). Since we ensured low expression concentration and chose high frame rates that result in high read noise, we chose to have high laser intensities to ensure that the fluctuation intensity was maximized for FCS, without significantly photobleaching the fast diffusing molecules seen in this study. Within the margins of error, the intensity in this range didn’t change the measured D [78].

### 4.6 SPIM-sFCCS Analysis

Autocorrelations and spatial cross-correlations on the acquired and simulated image stacks were performed using an ImageJ plugin Imaging FCS 1.52 modified for the fitting model given below. The intensity traces of the different pixels were autocorrelated, or traces from two specific pixels separated by a certain distance in x and y directions were cross-correlated over time *τ* as given by the following equation.

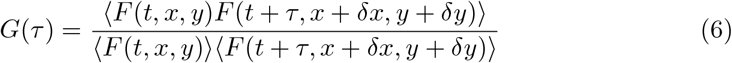

In the case of autocorrelations, *δx* and *δy* are 0. The diffusion coefficients were obtained by fitting the auto- and cross-correlation curves to a free 3D diffusion fitting model. The fitting equation derived in Wohland *et al*. (2010)[40] was modified to account for crosstalk between pixels from out-of-focus particles (5). This modifies the detection volume, which is illustrated in Fig. 2A. The modified cross-correlation model was thus

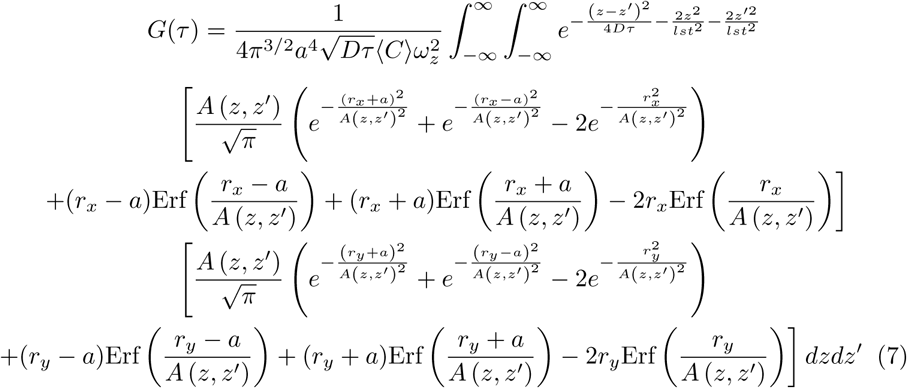

where

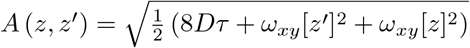

a = side length of a square pixel

D = the diffusion coefficient

*ω*_*z*_ = the 1*/e*^2^ radius of the light sheet thickness

*r*_*x*_ and *r*_*y*_ are distances between pixel pairs in x and y for which the cross-correlations are performed.

C = concentration; C is related to the number density in the pixel observation volume

V by C = N/V.

This equation was numerically integrated between –2*ω*_*z*_ and 2*ω*_*z*_ to capture almost all of the information for the Gaussian beam in the z-axis. The derivation for the above equation can be found in Appendix A.

The value for the pixel size (a) used for fitting was 400 nm, the value *ω*_*xy*_ used was 515 nm, and the value of *ω*_*z*_ was 1.13 *μm*.

The recorded data was checked for any movement and drift-related artifacts as described in Optimizing SPIM-sFCCS for *in vivo* measurements section by checking the forward and reverse fit results and including only the results where the distribution (Mean ± SD) of fit values overlapped.

The limited time resolution of the camera (β 1 ms) can result in the inaccurate fitting of auto-correlations capturing fast diffusion coefficients (60 *μm*^2^/s in Fig. 2 D.) Moreover, unlike in simulations, subtle background cell movements can cause low-frequency fluctuations in fluorescence intensity traces of the acquired data, which are unrelated to diffusion. These artifacts influence the longer lag time correlations, often resulting in correlation functions similar to a two-component ACF function, or can bias the single component fits to lower D values. Therefore, to reliably fit autocorrelations from acquired data, we binned pixels to create a large area for molecules to spend sufficient time to provide statistics for autocorrelations and only included the early lag times (*<* 100 ms) for fitting (Fig. 2E and for results in Fig. 5D).

All CCFs were fit assuming the presence of only a single diffusing component. Since pixel binning of at most 4 pixels was used, the CCFs over space were taken apart with 5, (2 *μm*), 9 (3.6 *μm*), 13 (5.2 *μm*) pixel separation distances chosen (Fig. 5D)

For diagonal channels (examples in Fig. 5A narrow channel, Supplementary Fig. 4F), a combination of x and y separations resulting in the equivalent separation distances as given above were used.

## Acknowledgements

We thank the NUS Centre for Bioimaging Sciences, SingaScope for providing microscope facility support and the Comparative Medicine (CM) fish facility for providing zebrafish care. The PMT-mApple plasmid used was a gift from Le Yao/Christoph Winkler. TW and TJC acknowledge funding from the Singapore Ministry of Education (MOE2016-T3-1-005). AVS and BS are supported by NUS Research Scholarships.

## Supplementary information

A supplementary file containing 2 appendix sections and 4 supplementary figures is included with this article.

## Declarations

### Ethics approval

All zebrafish experiments were performed following an approved protocol (IACUC #BR18-1023) by the Institutional Animal Care and Use Committee at the National University of Singapore.

### Data availability

Data can be provided upon reasonable request.

### Materials availability

Plasmids and maps can be provided on request.

### Code availability

ImFCS is a plugin in Fiji/ImageJ and is available publicly. Mathematica codes can be provided on request

### Author Contributions

TW, KS and TJC conceived the project. AVS and BS performed SPIM-FCS measurements. AVS and JS analyzed the data. AVS wrote the first draft, and all authors edited the final manuscript.

