## Supplementary Material for "Measuring morphogen transport over multiple spatial scales in live zebrafish embryos"

### Appendix A SPIM-FCS fit model for spatial cross-correalations

Here, we derive a SPIM-FCS model that accounts for the increased out-of-focus signal and, thus, the crosstalk between pixels in a light sheet system, as discussed in the main text.

The cross-correlation of the fluorescence fluctuations in two regions over time can be written as

$$G(\tau) = \frac{< F_1(t)F_2(t+\tau) >}{< F_1(t) >< F_2(t+\tau) >} \tag{A1}$$

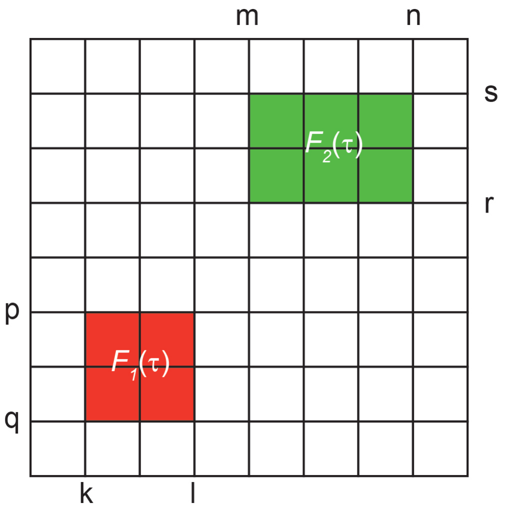


**Fig. A1 Illustration of the camera space:**
The red and green regions are the two areas being correlated in space

Where $F_i(t)$ is the fluorescence signal coming from region i = 1 or 2 (Fig. A1), $\tau$ is the time lag for which the correlation is calculated, and <> represents the time average.

The fluorescence intensity detected by a pixel will depend on the distribution of particles in the samples, $C(\vec{r}, t)$, the excitation intensity at the location of each particle, $I(\vec{r})$, which we assume to be constant in time, and the collection efficiency of the optical system for the particle location $\vec{r}$ in respect to the detection location $\vec{r}_0$, $CEF(\vec{r} - \vec{r}_0)$. We assume that the sample is much larger than the observation volume and thus is not explicitly taken into account in the derivation.

In SPIM-FCS, the light sheet thickness and intensity along the light sheet axis vary very slightly with position, and thus, we assume that the excitation intensity is constant in the xy-planes and depends only on the z-position (the optical axis of the detection objective) for a particular pixel. If one measures over an extent of the light sheet larger than the depth of focus, one can take account of the light sheet thickness by varying this thickness for different pixels. With these assumptions, we can write the intensity as

$$I(\overrightarrow{r}) = \frac{I_0}{(\frac{\pi}{2})^{\frac{1}{2}}\omega_z} e^{\frac{-2z^2}{\omega_z^2}} \tag{A2}$$

and the CEF as

$$CEF(\overrightarrow{r} - \overrightarrow{r}_0) = \frac{C_0 e^{\frac{-2((x-x_0)^2+(y-y_0)^2)}{\omega_{xy}[z-z_0]^2}}}{\frac{\pi}{2}\omega_{xy}[z-z_0]^2} \tag{A3}$$

where the spatial coordinate $\overrightarrow{r} = (x, y, z)$ describes the position of the emitting particle. The function $\omega_{xy} + |z|tan(\alpha)$ describes the increase of the image of out-of-focus molecules on the detector, with $\alpha$ being the half-collection angle of the objective. Note, that other models for the PSF can be easily integrated here. Our particular choice was governed by its simplicity and speed of calculation and its ability to fit the experimental data correctly, as tested by simulations (Fig. 2D). Eqs. A2 and A3 are normalized, so their integral results in $I_0$ or $C_0$, constants describing the intensity and CEF, taking into account the absolute excitation intensity and the properties of the fluorophore (extinction coefficient, quantum yield) and the detection properties of the optical system. The coordinates $\overrightarrow{r}_0 = (x_0, y_0, z_0)$ represent the position of the detection element. We assume that the camera's detector coincides with the image plane without any tilt, and thus we can set $z_0 = 0$,. The coordinates $(x_0, y_0)$ will range over a chosen pixel. Thus

$$CEF(\overrightarrow{r} - \overrightarrow{r}_0) = \frac{C_0 e^{\frac{-2((x-x_0)^2+(y-y_0)^2)}{\omega_{xy}[z]^2}}}{\frac{\pi}{2}\omega_{xy}[z]^2} \tag{A4}$$

For simplicity, in later calculations, we define the following function

$$W(\overrightarrow{r} - \overrightarrow{r}_0) = I(\overrightarrow{r})CEF(\overrightarrow{r} - \overrightarrow{r}_0) = \frac{I_0 C_0 e^{\frac{-2((x-x_0)^2+(y-y_0)^2)}{\omega_{xy}[z]^2}} e^{\frac{-2z^2}{\omega_z^2}}}{(\frac{\pi}{2})^{\frac{3}{2}}\omega_z\omega_{xy}[z]^2} \tag{A5}$$

With these definitions, we can describe the fluorescence intensities $F_i(t)$ seen by a pixel $i$ at position $(x_0, y_0)$ by

$$F_i(t) = \int_{x_0,y_0} \int_{-\infty}^{\infty} W(\overrightarrow{r} - \overrightarrow{r}_0) C(\overrightarrow{r}, t) d\overrightarrow{r} d\overrightarrow{r}_0 \tag{A6}$$

Here the integrals over $d\overrightarrow{r}_0$ range over the detection area of the pixel, which are described by the coordinates $(x_0 \in \{p, q\}, y_0 \in \{k, l\})$ for pixel 1 and $(x_0 \in \{m, n\}, y_0 \in \{r, s\})$ for pixel 2 (Fig. A1). The integral over $d\overrightarrow{r}$ ranges over the entire space, summing the fluorescence contribution to a particular pixel of all particles. Substituting these equations into the equation for the correlation function (Eq. A1) results in

$$G(\tau) = \frac{\int_p^q \int_k^l \int_r^s \int_m^n \int_{-\infty}^{\infty} \int_{-\infty}^{\infty} W(\overrightarrow{r} - \overrightarrow{r}_0) W(\overrightarrow{r}' - \overrightarrow{r}'_0) \langle C(\overrightarrow{r}, t) C(\overrightarrow{r}', t+\tau) \rangle d\overrightarrow{r} d\overrightarrow{r}' dx_0 dy_0 dx'_0 dy'_0}{(\int_p^q \int_k^l \int_{-\infty}^{\infty} W(\overrightarrow{r} - \overrightarrow{r}_0) \langle C \rangle d\overrightarrow{r} dx_0 dy_0)(\int_r^s \int_m^n \int_{-\infty}^{\infty} W(\overrightarrow{r} - \overrightarrow{r}_0) \langle C \rangle d\overrightarrow{r} dx_0 dy_0)} \tag{A7}$$

Here, we have made the integration over the pixel areas ($x_0 \in \{p, q\}, y_0 \in \{k, l\}$) and ($x_0 \in \{m, n\}, y_0 \in \{r, s\}$) explicit. Note that the time average is taken only over the concentrations, as all other values are constant. The resulting expression $\langle C(x, y, z, t) C(x', y', z', t+\tau) \rangle$ describes the probability that a particle at position $\overrightarrow{r}$ at time $t$ can be found at position $\overrightarrow{r}'$ at time $t + \tau$, and is known as the diffusion propagator [1], [2], [3].

$$\langle C(\overrightarrow{r}, t) C(\overrightarrow{r}', t+\tau) \rangle = \langle C \rangle \frac{e^{\frac{-(\overrightarrow{r} - \overrightarrow{r}')^2}{4D\tau}}}{(4\pi D\tau)^{\frac{3}{2}}} \tag{A8}$$

This results in

$$G(\tau) = \frac{\int_p^q \int_k^l \int_r^s \int_m^n \int_{-\infty}^{\infty} \int_{-\infty}^{\infty} W(\overrightarrow{r} - \overrightarrow{r}_0) W(\overrightarrow{r}' - \overrightarrow{r}'_0) \frac{e^{\frac{-(r - \overrightarrow{r}')^2}{4D\tau}}}{(4\pi D\tau)^{\frac{3}{2}}} d\overrightarrow{r} d\overrightarrow{r}' dx_0 dy_0 dx'_0 dy'_0}{\langle C \rangle \int_p^q \int_k^l \int_{-\infty}^{\infty} W(\overrightarrow{r} - \overrightarrow{r}_0) d\overrightarrow{r} dx_0 dy_0 \int_r^s \int_m^n \int_{-\infty}^{\infty} W(\overrightarrow{r} - \overrightarrow{r}_0) d\overrightarrow{r} dx_0 dy_0} \tag{A9}$$

This function can be integrated for x and y dimensions [4]. The integrals in the denominator result in

$$\int_p^q \int_k^l \int_{-\infty}^{\infty} W(\overrightarrow{r} - \overrightarrow{r}_0) d\overrightarrow{r} dx_0 dy_0 = I_0 C_0 (l - k)(q - p) \tag{A10}$$

$$\int_r^s \int_m^n \int_{-\infty}^{\infty} W(\overrightarrow{r} - \overrightarrow{r}_0) d\overrightarrow{r} dx_0 dy_0 = I_0 C_0 (n - m)(s - r) \tag{A11}$$

Equation A9 now simplifies to

$$G(\tau) = \frac{1}{I_0^2 C_0^2 \langle C \rangle (l-k)(q-p)(n-m)(s-r)}$$
$$\int_p^q \int_k^l \int_r^s \int_m^n \int_{-\infty}^{\infty} \int_{-\infty}^{\infty} W(\overrightarrow{r} - \overrightarrow{r}_0) W(\overrightarrow{r}' - \overrightarrow{r}_0)' \frac{e^{\frac{-(r - \overrightarrow{r}')^2}{4D\tau}}}{(4\pi D\tau)^{\frac{3}{2}}} d\overrightarrow{r} d\overrightarrow{r}' dx_0 dy_0 dx'_0 dy'_0 \tag{A12}$$

Expanding Eq. A12 using the relation defined in Eq. A5, we obtain the following result:

$$G(\tau) = \frac{1}{\left(\pi^{3/2}(D\tau)\right)^{3/2}(l-k)(n-m)(q-p)(s-r)\langle C\rangle}$$
$$\int_r^s\int_m^n\int_p^q\int_k^l\int_{-\infty}^{\infty}\int_{-\infty}^{\infty}\int_{-\infty}^{\infty}\int_{-\infty}^{\infty}\int_{-\infty}^{\infty}\int_{-\infty}^{\infty}\frac{1}{\left(\pi^{3/2}\omega_z\right)\omega_{xy}[z]^2}$$
$$\frac{1}{\left(\pi^{3/2}\omega_z\right)\omega_{xy}[z']^2}\exp\left(-\frac{(x-x')^2}{4D\tau}-\frac{(y-y')^2}{4D\tau}-\frac{(z-z')^2}{4D\tau}\right)$$
$$\exp\left(-\frac{2\left(x-x_0\right)^2}{\omega_{xy}[z]^2}-\frac{2\left(y-y_0\right)^2}{\omega_{xy}[z]^2}-\frac{2z^2}{\omega_z^2}\right)$$
$$\exp\left(-\frac{2\left(x'-x_0'\right)^2}{\omega_{xy}[z']^2}-\frac{2\left(y'-y_0'\right)^2}{\omega_{xy}[z']^2}-\frac{2\left(z'\right)^2}{\omega_z^2}\right)dxdx_0dydy_0dzdx'dx_0'dy'dy_0'dz' \quad \text{(A13)}$$

This integral can be grouped into x and y components. Note that they are still a function of z. The integration over $x$,$x_0$,$y$ and $y_0$ can be performed first. The following shows the grouping into x and y components:

$$xcomponent : \int_m^n\int_k^l\int_{-\infty}^{\infty}\int_{-\infty}^{\infty}\exp\left(-\frac{(x-x')^2}{4D\tau}-\frac{2\left(x'-x_0'\right)^2}{\omega_{xy}[z']^2}-\frac{2\left(x-x_0\right)^2}{\omega_{xy}[z]^2}\right) dxdx'dx_0dx_0' \quad \text{(A14)}$$

$$ycomponent : \int_r^s\int_p^q\int_{-\infty}^{\infty}\int_{-\infty}^{\infty}\exp\left(-\frac{(y-y')^2}{4D\tau}-\frac{2\left(y'-y_0'\right)^2}{\omega_{xy}[z']^2}-\frac{2\left(y-y_0\right)^2}{\omega_{xy}[z]^2}\right) dydy'dy_0dy_0' \quad \text{(A15)}$$

$zcomponent :$

$$\int_{-\infty}^{\infty}\int_{-\infty}^{\infty}\frac{1}{\left(\pi^{3/2}\omega_z\right)^2\omega_{xy}[z]^2\omega_{xy}[z']^2}\exp\left(-\frac{(z-z')^2}{4D\tau}-\frac{2z^2}{\omega_z^2}-\frac{2\left(z'\right)^2}{\omega_z^2}\right)dzdz' \quad \text{(A16)}$$
$$denominator : \frac{1}{\left(\pi^{3/2}(D\tau)\right)^{3/2}(l-k)(n-m)(q-p)(s-r)\langle C\rangle} \quad \text{(A17)}$$

Taking the different components together, we have

$$G(\tau) = (xcomponent)(ycomponent)(zcomponent)(denominator)$$

The solution for the integral in Eqs. A14 and A15 can be obtained analytically as shown in Sankaran et al. (2009)[5]. The result for Eq. A13 then is:

$$\frac{1}{2}\pi^{3/2}\sqrt{D\tau}\omega_{xy}[z]\omega_{xy}[z']\left(\frac{\sqrt{8D\tau+\omega_{xy}[z']^2+\omega_{xy}[z]^2}}{\sqrt{2\pi}}\right)$$

$$\left(\exp\left(-\frac{2(k-n)^2}{8D\tau+\omega_{xy}[z']^2+\omega_{xy}[z]^2}\right)-\exp\left(-\frac{2(k-m)^2}{8D\tau+\omega_{xy}[z']^2+\omega_{xy}[z]^2}\right)\right.$$

$$\left.+\exp\left(-\frac{2(l-m)^2}{8D\tau+\omega_{xy}[z']^2+\omega_{xy}[z]^2}\right)-\exp\left(-\frac{2(l-n)^2}{8D\tau+\omega_{xy}[z']^2+\omega_{xy}[z]^2}\right)\right)$$

$$-(m-k)\mathrm{erf}\left(\frac{m-k}{\sqrt{\frac{1}{2}\left(8D\tau+\omega_{xy}[z']^2+\omega_{xy}[z]^2\right)}}\right)$$

$$+(m-l)\mathrm{erf}\left(\frac{m-l}{\sqrt{\frac{1}{2}\left(8D\tau+\omega_{xy}[z']^2+\omega_{xy}[z]^2\right)}}\right)$$

$$+(n-k)\mathrm{erf}\left(\frac{n-k}{\sqrt{\frac{1}{2}\left(8D\tau+\omega_{xy}[z']^2+\omega_{xy}[z]^2\right)}}\right)$$

$$-(n-l)\mathrm{erf}\left(\frac{n-l}{\sqrt{\frac{1}{2}\left(8D\tau+\omega_{xy}[z']^2+\omega_{xy}[z]^2\right)}}\right) \tag{A18}$$

The terms in the x and y solutions can be conveniently simplified through the following substitution:

$$A\left(z,z'\right)=\sqrt{\frac{1}{2}\left(8D\tau+\omega_{xy}[z']^2+\omega_{xy}[z]^2\right)} \tag{A19}$$

Using the results from Eq. A18 and A19, the final expression of $G(\tau)$ as a function of $z$ and $z'$ is

$$G(\tau)=\frac{1}{4\pi^{3/2}\sqrt{D\tau}(l-k)(n-m)(q-p)(s-r)\langle C\rangle\omega_z^2}$$

$$\int_{-\infty}^{\infty}\int_{-\infty}^{\infty}e^{-\frac{(z-z')^2}{4D\tau}-\frac{2z^2}{\omega_z^2}-\frac{2z'^2}{lst^2}}$$

$$\left[\frac{A\left(z,z'\right)}{\sqrt{\pi}}\left(-e^{-\frac{(k-m)^2}{A(z,z')^2}}+e^{-\frac{(k-n)^2}{A(z,z')^2}}+e^{-\frac{(l-m)^2}{A(z,z')^2}}-e^{-\frac{(l-n)^2}{A(z,z')^2}}\right)\right.$$

$$-(m-k)\mathrm{Erf}\left(\frac{m-k}{A\left(z,z'\right)}\right)+(m-l)\mathrm{Erf}\left(\frac{m-l}{A\left(z,z'\right)}\right)$$

$$\left.+(n-k)\mathrm{Erf}\left(\frac{n-k}{A\left(z,z'\right)}\right)-(n-l)\mathrm{Erf}\left(\frac{n-l}{A\left(z,z'\right)}\right)\right]$$

$$\left[\frac{A\left(z,z'\right)}{\sqrt{\pi}}\left(-e^{-\frac{2(p-r)^2}{A(z,z')^2}}+e^{-\frac{2(p-s)^2}{A(z,z')^2}}+e^{-\frac{(q-r)^2}{A(z,z')^2}}-e^{-\frac{(q-s)^2}{A(z,z')^2}}\right)\right.$$

$$+(p-r)\mathrm{Erf}\left(\frac{r-p}{A\left(z,z'\right)}\right)+(r-q)\mathrm{Erf}\left(\frac{r-q}{A\left(z,z'\right)}\right)$$

$$+(s-p)\text{Erf}\left(\frac{s-p}{A\left(z,z'\right)}\right)+(q-s)\text{Erf}\left(\frac{s-q}{A\left(z,z'\right)}\right)\Bigg]\,dzdz' \tag{A20}$$

Substituting the values of k=0, l=a, m=$r_x$, n=a+$r_x$, p=0, q=a, r=$r_y$, s=a+$r_y$, the generic function for cross-correlation is as follows.

$$G(\tau)=\frac{1}{4\pi^{3/2}a^4\sqrt{D\tau}\langle C\rangle\omega_z^2}\int_{-\infty}^{\infty}\int_{-\infty}^{\infty}e^{-\frac{(z-z')^2}{4D\tau}-\frac{2z^2}{\omega_z^2}-\frac{2z'^2}{lst^2}}$$

$$\left[\frac{A\left(z,z'\right)}{\sqrt{\pi}}\left(e^{-\frac{(r_x+a)^2}{A(z,z')^2}}+e^{-\frac{(r_x-a)^2}{A(z,z')^2}}-2e^{-\frac{r_x^2}{A(z,z')^2}}\right)\right.$$

$$\left.+(r_x-a)\text{Erf}\left(\frac{r_x-a}{A\left(z,z'\right)}\right)+(r_x+a)\text{Erf}\left(\frac{r_x+a}{A\left(z,z'\right)}\right)-2r_x\text{Erf}\left(\frac{r_x}{A\left(z,z'\right)}\right)\right]$$

$$\left[\frac{A\left(z,z'\right)}{\sqrt{\pi}}\left(e^{-\frac{(r_y+a)^2}{A(z,z')^2}}+e^{-\frac{(r_y-a)^2}{A(z,z')^2}}-2e^{-\frac{r_y^2}{A(z,z')^2}}\right)\right.$$

$$\left.+(r_y-a)\text{Erf}\left(\frac{r_y-a}{A\left(z,z'\right)}\right)+(r_y+a)\text{Erf}\left(\frac{r_y+a}{A\left(z,z'\right)}\right)-2r_y\text{Erf}\left(\frac{r_y}{A\left(z,z'\right)}\right)\right]dzdz' \tag{A21}$$

However, as the z dimension is not independent of the x and y components, the integral has no analytical solution. Therefore, to fit data, we numerically integrate the above equation between $-2\omega_z$ to $2\omega_z$ as >99 % of the extent of the light sheet is captured within $2\omega_z$ in either direction of the z-axis.

The precision of Eq. A21, depends on the numerator, which is dependent on $z$ and $z'$, and thus the number of segments in the numerical integration; and it depends on the denominator, which has a term $\sqrt{D\tau}$. As $\sqrt{D\tau}$ will decrease with D and $\tau$, the value of Eq. A21 will start diverging for short lag times and low diffusion coefficients if the numerical integration in the nominator is not sufficiently accurate (Fig. A2). Therefore, we increase the number of segments in the numerical integration with decreasing diffusion coefficients to obtain the desired accuracy.

Empirically, for our time resolution of $\tau \geq 1$ ms and light sheet thickness $\omega_z = 1.13$ $\mu m$, we found that $\delta\omega_z = \omega_z\ /5$ (20 integration segments) captures D $\geq 5$ $\mu m^2/s$ correctly but fails for smaller D (Fig. A2A, B). $\delta\omega_z = \omega_z/20$ (80 integration segments) is adequate for D $\geq 0.5$ $\mu m^2/s$; and $\delta\omega_z = \omega_z/30$ (120 integration segments) is sufficient for D $\geq 0.3$ $\mu m^2/s$ (Fig. A2C, D).

Since increasing the segment number also increases calculation costs and fitting time, we used a medium segment number of 80, i.e. $\delta\omega_z = \omega_z\ /20$, as this covered most of the experimental values of D $> 0.5$ $\mu m^2/s$. If smaller D values are to be measured, the segment number needs to be adjusted.

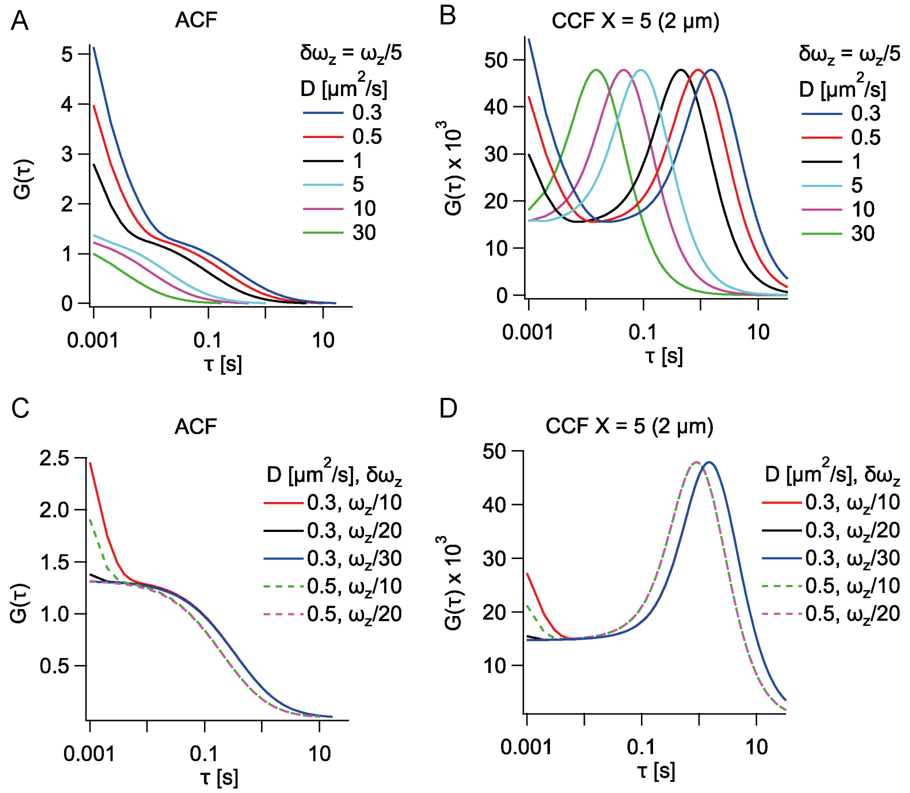


**Fig. A2 Comparison of the ACFs and CCFs for different segment sizes used for the numerical integration:**
A. Plots of ACFs calculated for different D values for a large $\delta\omega_z$ segment size of $\omega_z/5$. Note how the ACF diverges at short lag times for D $\leq$ 1 $\mu m^2$/s.
B. Plots of CCFs calculated for different D values for a large segment size of $\omega_z$ /5. Similar to the ACF, the CCF diverges at short lag times for D $\leq$ 1 $\mu m^2$/s.
C. Plots of ACFs calculated for lower D values of 0.3 and 0.5 $\mu m^2$/s and smaller $\delta\omega_z$ segment sizes. With decreasing $\delta\omega_z$ segment size, the divergence is reduced for the short lag time values, resulting in an improvement in fitting accuracy .
D. Plots of CCFs calculated for lower D values of 0.3 and 0.5 $\mu m^2$/s and smaller $\delta\omega_z$ segment sizes. Similar to ACFs, with decreasing $\delta\omega_z$ segment size, the accuracy on the short lag time value improves for CCFs.

#### Appendix B BMP sfGFP $D_{eff}$ measured at 3.6 $\mu m$ separation distance.

| Channel Type | $D_{eff}(\mu m^2/s)$ | $N_{ccf}$ |
|---|---|---|
| Narrow | 17.3 ± 3.8 | 20 |
| (< 2.5 $\mu m$) | 10.3 ± 3.9 | 20 |
| | 17.5 ± 3.8 | 32 |
| $D_{avg}$ = 15.0 ± 4.6 ($N_{emb}$ = 3) | | |
| Wide | 57.3 ± 11.6 | 132 |
| (> 4 $\mu m$) | 50.1 ± 7.1 | 54 |
| | 58.0 ± 19.6 | 132 |
| | 58.4 ± 18.0 | 36 |
| | 56.5 ± 21.1 | 50 |
| $D_{avg}$ = 56.1 ± 16.3 ($N_{emb}$ = 4) | | |

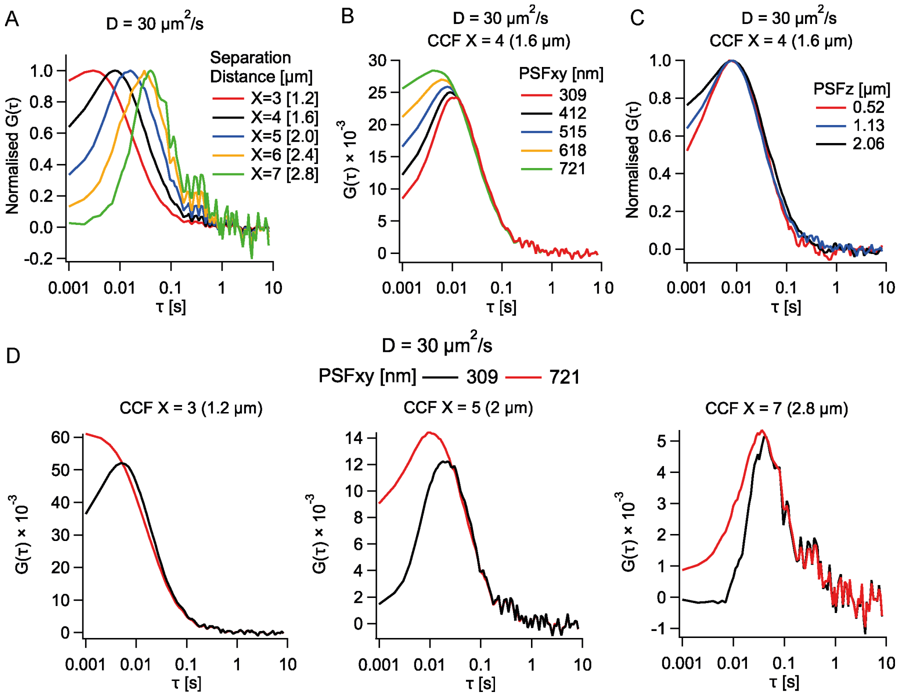


**Supplementary Fig. 1: Influence of signal cross-talk on the CCF:**
A. Comparison of the raw average cross-correlation function from multiple correlations for different separation distances across space. The component of pseudo-autocorrelation component extends almost over 6 pixels (2.4 $\mu m$).
B. Comparison of the raw average cross-correlation function from multiple correlations for different $\mathrm{PSF}_{xy}$ values simulated. With increasing $\mathrm{PSF}_{xy}$ value, the contribution of the crosstalk component increases.
C. Comparison of the raw average cross-correlation function from multiple correlations for different light sheet thickness values simulated. With increasing light sheet thickness ($\mathrm{PSF}_z$) value, there is a slight increase in the cross talk component.
D. Comparison of the cross-correlation functions of two different $\mathrm{PSF}_{xy}$ values across space. Note that with increasing separation distance, the difference between the cross-correlation functions corresponding to the two different PSF values reduces.

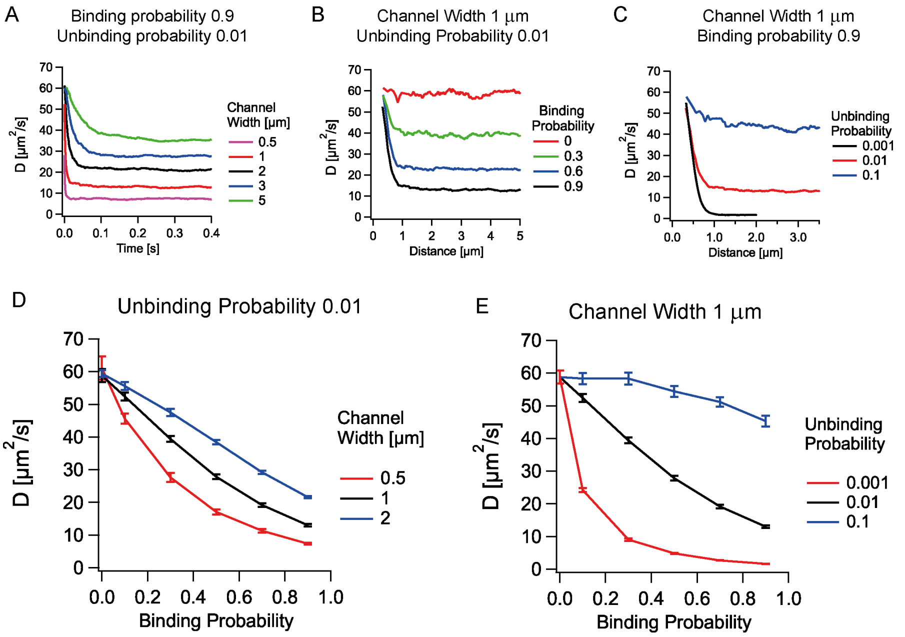


**Supplementary Fig. 2: Influence of different parameters on the diffusion of particles in the intermembrane-like spaces between cells:**

A. Plot of the transition of effective diffusion coefficients over time for different channel widths. This result for time is converted to the distance traversed and plotted in Fig. 3B.

B. Plot of the transition of effective diffusion coefficients over distance for different binding probabilities. The effective diffusion coefficient decreases with increasing binding probability, a proxy for receptor concentration on the membrane.

C. Plot of the transition of effective diffusion coefficients over distance for different channel widths. The effective diffusion coefficient decreases with decreasing unbinding probability. Low unbinding probability is indicative of a high ligand-receptor binding affinity .

D. Plot of the steady-state effective diffusion coefficient reached vs. the binding probability set for different narrow channel widths. Error bars are the SD of values from 3 independent simulation seeds. The effective diffusion coefficients are lower for smaller channels and higher binding probabilities.

E. Plot of the steady-state effective diffusion coefficient reached vs. the binding probability set for different unbinding probabilities. Error bars are the SD of values from 3 independent simulation seeds. The effective diffusion coefficients decrease with the lowering of the unbinding probability values.

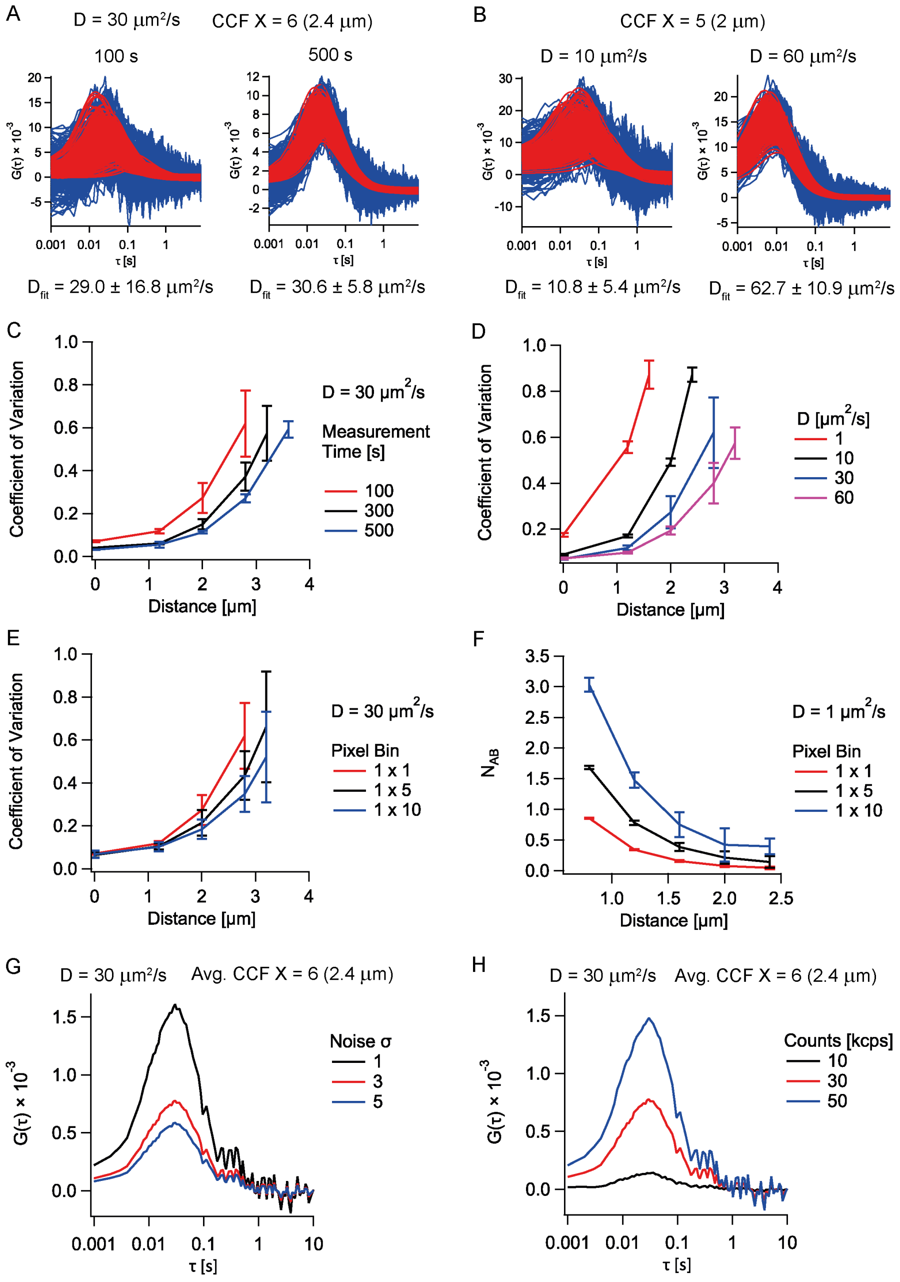


**Supplementary Fig. 3: Effects of physical parameters on CCF signal:**
A. Comparison of the raw cross-correlation function (blue) obtained and their corresponding fits (red) for different measurement times (simulated data). With increasing measurement time, the quality of correlations improves, resulting in better fits and low SDs in the $D_{fit}$ results.

**Previous page caption continued:**
B. Comparison of the raw cross-correlation function (blue) obtained and their corresponding fits (red) for different Ds (simulated data). With increasing D value to be measured, the quality of correlations improves, resulting in better fits and low SDs in the $D_{fit}$ results.
C. Coefficient of variation (COV = SD/Mean value) among individual pairs of cross-correlation fits over space for fit data obtained from 3D free diffusion simulations of different measurement times. Error bars: SD over three independent simulation sets. Note the lower COV and longer range for higher measurement times.
D. COV among individual pairs of cross-correlation fits over space for fit data obtained from simulations of different diffusion coefficients. Error bars: SD over three independent simulation seeds. Note the comparatively lower COV and longer range for higher D values.
E. COV among individual pairs of cross-correlation fits over space for fit data obtained from simulations while adopting different spatial pixel bins. Error bars: SD over three independent simulation seeds. Note the comparatively lower COV and longer range for higher bin values.
F. Comparison between different bin sizes on how many transit particles are captured between two points in space (simulated data). Error bars: SD over three independent simulation seeds. Larger bin sizes allow for more particles to be captured.
G. Average cross-correlation curves for different background Gaussian noise values (simulated data). Note the decrease in CCF amplitude with increasing noise levels.
H. Average cross-correlation curves for different counts emitted by a diffusing particle (simulated data). Note the increase in CCF amplitude with increasing counts emitted.

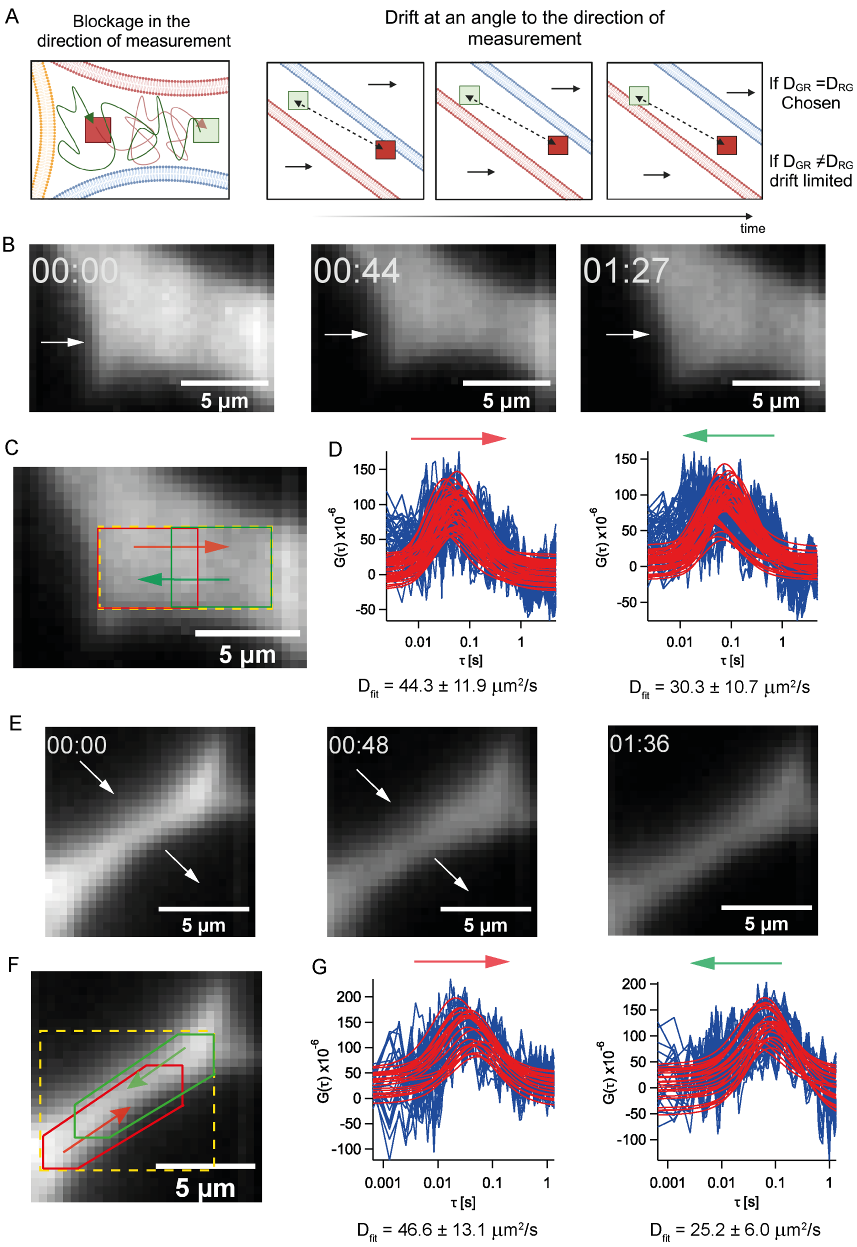


**Supplementary Fig. 4: Influence of cell movements on SPIM-sFCCS:**
A. Illustration of the effects of barriers and cell movements/drifts on the $D_{eff}$ measured over space. Barriers to diffusion cause reflections and contribute twice to transit dynamics while drifts can add or subtract to the diffusion coefficient of molecules depending on the direction measured. Accordingly, only cases where the forward and backward components of the measured D values are equal should be chosen.

**Previous page caption continued:**
B. Example of a Sqt-EGFP measurement to show how cells move over time and push molecules in the cell spaces in a tissue.
C. The region of interest to measure the effective diffusion coefficients for the channel.
D. Spatial cross-correlation data obtained in the forward and reverse directions (raw correlations: blue, fits: red) from data in C. The measurement performed against the direction of cell movements yielded slower-than-usual effective diffusion coefficient values. The unequal values indicate the influence of cell movements to transport or block the diffusion of molecules over space.
E. Example of a Sqt-EGFP measurement to show channel movement and changing of channel shape over time.
F. The region of interest to measure the effective diffusion coefficients for the channel.
G. Spatial cross-correlation data obtained in the forward and reverse directions (raw correlations: blue, fits: red) from data in F. The unequal values for the two cases indicate the influence of cell movements.
